# Decoding human sperm signalling: Phosphoproteomic discovery of kinases governing fertilization competency

**DOI:** 10.64898/2026.06.22.733301

**Authors:** Nathan D Burke, Amanda L Anderson, John E Schjenken, Shaun D Roman, Hanah M Hart, Heather C Murray, Kasey Miller, Georgia E Blackley, R John Aitken, David A Skerrett-Byrne, Brett Nixon, Elizabeth G Bromfield

## Abstract

Capacitation, the process whereby sperm gain the functional competence to fertilize an egg in the absence of de novo transcription and translation, is orchestrated by a hierarchy of kinases driving the phosphorylation of sperm proteins. While increased phosphorylation, in particular tyrosine phosphorylation, is a revered hallmark of fertilization competency in our species, only a limited repository of phosphorylated substrates and kinases have ever been reported from human sperm.

To broaden therapeutic targets for sperm targeted contraceptives and infertility therapies, we adapted a contemporary phosphoproteomic technique termed EasyPhos to generate bespoke methodology for the investigation of human sperm signalling. This approach yielded high depth phosphoproteomes of non-capacitated and capacitated human spermatozoa with in silico investigation of the phosphosites revealing 52 kinases with previously uncharacterized roles in sperm capacitation. Investigating the function of the putative sperm capacitation kinases identified yielded several kinases with novel roles in the regulation of sperm function. Of particular interest, polo like kinase 1 (PLK1) inhibition significantly reduced progressive sperm motility, attenuated capacitation-associated tyrosine phosphorylation and reduced the sperm acrosome reaction, an essential step to achieve fertilization. These findings reveal extensive phosphoproteome remodelling during human sperm capacitation, expanding the landscape of molecular targets for fertility control.

## INTRODUCTION

Morphologically mature spermatozoa are intricately fashioned by mitotic expansion of spermatogonia followed by a series of nuclear reduction events (meiosis), organelle remodeling, the fabrication of sperm-specific organelles (e.g. acrosome, manchette, and fibrous sheath) and cytoplasmic shedding (spermiogenesis) ^1–3^. Despite the innate complexity of spermatogenesis, morphologically mature testicular sperm remain impotent and unable to achieve fertilization and must therefore complete two distinct sequential processes of functional maturation before realizing their potential to fertilize an egg ^4^.

Epididymal transit, the first of these sequential maturation processes, confers fertilization potential via macromolecular exchange with the luminal milieu of the epididymis ^5–9^. Recent proteomic studies in mice have detailed the refinement of the sperm proteome during epididymal transit, including a halving of the sperm protein complement in favor of an enrichment of proteins that are crucial for fertilization^10^. Phosphoproteomic inquiry into the functional modification of the mouse sperm proteome throughout epididymal transit has demonstrated widespread remodeling of phosphorylation signatures ^11,12^. Specifically, Skerrett-Byrne *et al.* ^12^ report that 86% of altered phosphorylation events in mouse sperm occur during epididymal transit. Indeed, after completing epididymal transit, sperm are equipped with a curated inventory of proteins necessary to achieve fertilization, however, these cells still remain naïve to the egg, a quandary that was prohibitive to early attempts to develop *in vitro* fertilization in mammals. Independently, Austin ^13^ and Chang ^14^ revealed the requirement for sperm to take residency in the female reproductive tract (later termed capacitation ^15^), as the ultimate maturational milestone for sperm to gain the functional competence to fertilize an egg.

In eutherian mammals, sperm capacitation commences after insemination with the sequestration of sterols (including cholesterol and desmosterol) from the sperm plasma membrane which, *in vivo*, is facilitated by extracellular constituents including proteinaceous acceptor molecules such as albumin, high-density lipoproteins and apolipoproteins (e.g. APOA-1) ^16^. The corresponding increase in membrane permeability promotes an influx of bicarbonate ions and subsequent activation of Na^2^/H^+^ exchangers ^17^, and other ion channels ^18^, and ultimately precipitates an increased intracellular pH (intracellular alkalinization). This, in turn, gives rise to a microenvironment in which HCO ^-^ ions activate soluble adenylyl cyclase (sAC) production of cyclic adenosine 3^’^,5^’^ monophosphate (cAMP) ^19,20^. While a foray of other signaling events occur concomitantly (as reviewed by Gervasi and Visconti ^21^), including membrane hyperpolarization and increased intracellular Ca^2+^ ^22,23^, cAMP activates protein kinase A (PKA), a master orchestrator of the phosphorylation signaling cascade underpinning sperm capacitation ^24,25^. Remarkably, this fastidious cascade of post translational modifications activates and positions the spermatozoon’s previously quiescent fertilization machinery, in the absence of transcription and translation. These activation events enable hyperactivation of motility needed for oocyte penetration ^26^, sperm surface remodeling events ^27–29^, and sensitizes sperm to the physiological agonists of the female reproductive tract (e.g. progesterone and zona pellucida (ZP) glycoproteins) necessary to undergo an exocytotic event termed the acrosome reaction ^30^. Finally, capacitation facilitates the activation and positioning of sperm-egg recognition candidates (e.g. IZUMO1, TMEM95, and SPACA6) ^31,32^ in the now actualized sperm cell (as reviewed by Aitken and Nixon ^33^).

Despite long-held knowledge that capacitation increases protein phosphorylation, in particular protein tyrosine phosphorylation, an established phenomenon and typical hallmark of capacitation ^34,35^. However, only a limited repository of phosphorylated substrates ^36–39^ and kinases ^40–45^ have been characterized from capacitated sperm, especially in humans. This limited knowledge of sperm capacitation is currently prohibitive to the effective development of sperm-targeted contraceptives focused on preventing sperm maturation ^46^ and a limiting factor for the development of novel treatments for sperm-dysfunction, a major cause of infertility. The potential for proteomics to investigate both the molecular composition of sperm and the post translational modifications that regulate their function is well appreciated ^47–52^. In particular, Serrano *et al.* ^49^ note that, despite the advances in the field of phosphoproteomics, these technologies have been scarcely used for investigating human sperm and suggest that phosphoproteomics has not yet been applied to its full potential in studying male fertility. Despite this, several studies have made positive headway into investigating the role of protein phosphorylation in regulating spermatogenesis and cell survival in the human testes ^53^. Studies have also focused on the role of the phosphoproteome in human sperm motility ^54^, with comparisons also made between normozoospermic and asthenozoospermic men ^55,56^, fertile and infertile men ^57^, fresh and cryopreserved sperm ^58^, and non-capacitated and capacitated sperm ^36,37,59^. In this series of experiments, we employed new phosphoproteomic methodology ^60^ and *in silico* analyses ^61,62^ to gain a high-depth molecular map of phospho-driven control of sperm function. Building on this curated dataset of phosphorylation events, we conducted thorough kinase inhibition studies to examine whether the manipulation of the specific kinases revealed by our work show merit for the controlled modulation of sperm capacitation. These results reveal a previously unappreciated role for several kinases including polo like kinase 1 (PLK1) during capacitation in mature human spermatozoa.

## RESULTS

### Characterization of the human sperm phosphoproteome

To produce the human sperm phosphoproteome described in this study, we performed phosphopeptide enrichment and high-resolution tandem mass spectrometry (nLC-MS-MS) on ejaculated human sperm cells in a naïve, non-capacitated state and following the induction of *in vitro* capacitation. To ensure high quality sample preparation, viability and total motility were assessed across all sperm samples prior to mass spectrometry (Figure S1A) revealing equivalent levels of motility and viability in both populations and consistency across all biological replicates. To validate each sample’s capacitation status prior to their use for mass spectrometry, immunoblotting and immunolabeling with anti-phosphotyrosine, -phosphoserine and - phosphothreonine antibodies was performed on all sperm samples (Figure S1B, D, & E). As expected from capacitated human sperm ^37,38,63–65^, band densitometry and labeling pattern counts (Figure S1C and F respectively) demonstrated significantly increased protein tyrosine phosphorylation compared to their non-capacitated sperm counterparts. Phosphothreonine labeling of the flagellum was detected in both non-capacitated and capacitated sperm, however, in a similar trend to phosphotyrosine, phospho-threonine labeling was more intense in capacitated sperm. Accordingly, the tail labeling counts for phosphothreonine were performed on the basis of dim compared to bright staining (Figure S1 F), with representative population images included in Figure S1E depicting this trend.

Following nLC-MS-MS, an assessment of the correlation between biological replicates was performed demonstrating high similarity, with average Pearson correlations of 0.9 for non-capacitated samples and 0.86 for capacitated samples, respectively (Figure 1A). Phosphoproteomic profiling (false discovery rate [FDR] ≤ 0.01) revealed 825 and 840 phosphoproteins in non-capacitated (NC) and capacitated (CAP) human sperm (presented as NC and CAP in figures), respectively (Figure 1B), totaling 843 proteins (Table S1) with an overall phosphopeptide enrichment efficiency of 82.06%. Non-capacitated sperm yielded 1,719 phosphopeptides, with an average of 2.133 peptides per protein, while capacitated human sperm produced 1,765 phosphopeptides, with an average of 2.138 peptides per protein (Figure 1B). Principal component analysis (PCA) revealed that non-capacitated and capacitated human sperm exhibited discernible differences in their phosphoproteomes that were conserved across the five biological replicates. This indicated high reproducibility between samples, with two components accounting for 73.2% of the observed variance (Figure 1C).

**Figure 1:**
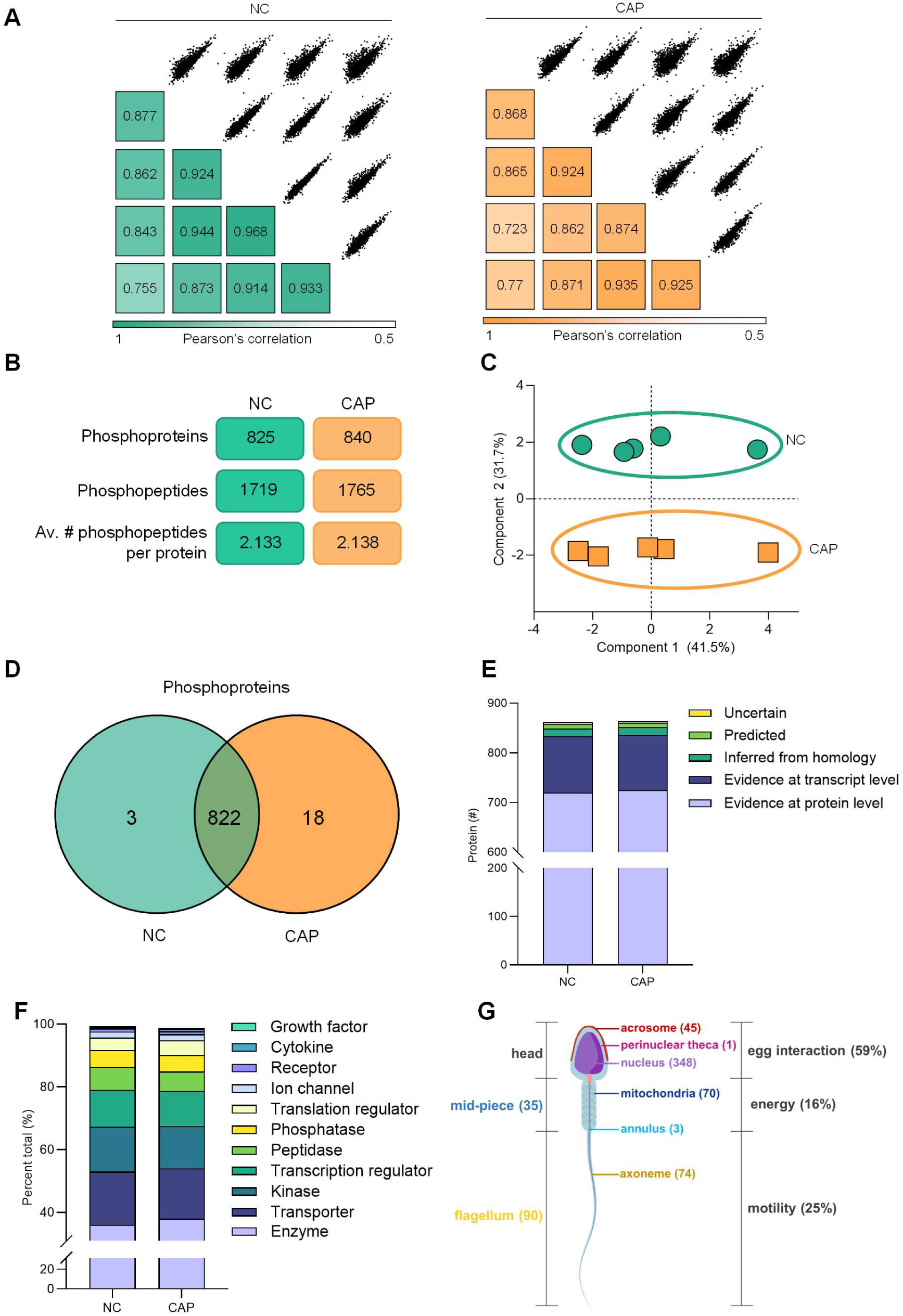
Characterization of the human sperm phosphoproteome. (A) Pearson correlation plots of biological replicates for non-capacitated (green) and capacitated (orange) human sperm populations. (B) The total number of phosphoproteins identified within each populations phosphoproteomic signature, non-capacitated (825, green), and capacitated (840, orange), along with the total number of peptides and average number of phosphopeptides/protein per population. (C) Principal component analysis (PCA) of the phosphoproteome of each sperm cell population with their respective biological replicates; non-capacitated (green) and capacitated (orange). (D) Venn diagram depicting the unique and common phosphoproteins between non-capacitated (NC) and capacitated (CAP) human sperm cells. (E) UniProt annotation of level of evidence for protein existence. (F) Classification of protein types by Ingenuity Pathway Analysis represented as a proportion of all proteins identified. (G) Phosphoprotein localization derived from UniProt Gene Ontology, note acrosome, perinuclear theca and nucleus regions of sperm are categorized as “egg interaction” regions, mid-piece and mitochondria are classified as “energy” producing regions, and the flagellum, anulus and axoneme are categorized as “motility” associated regions. A-G present in-silico proteomic analysis on 5 biological samples.

Direct comparison of the two sperm populations demonstrated a shared proteomic signature of 822 phosphoproteins (Figure 1D). Three proteins were uniquely detected in non-capacitated sperm compared to 18 uniquely detected in capacitated samples (Table S1). Assessment of the curated evidence (UniProt) for the existence of each protein identified revealed 148 proteins with no previous evidence at the protein level (Figure 1E and Table S1), consisting of 119 proteins with only transcript evidence (13.2%), 16 inferred homology (1.8%), and 9 predicted (0.1%). Compositional analysis of phosphoprotein signatures revealed that the human sperm phosphoproteome broadly consists of enzymes (37%), kinases (13.9%), phosphatases (5.2%), and ion channels (2%), among other receptors and immune proteins (Figure 1F and Table S1). Additional *in silico* analysis using UniProt Gene Ontology GO) annotations mapped 79% of the identified phosphoproteins (666) to a specific sperm cell domain or organelle (i.e. acrosome, perinuclear theca, nucleus, mitochondria, mid-piece, annulus, axoneme, and flagellum; mid-piece and flagellum occurred concomitantly with terms including connecting piece, principal piece, end piece, and fibrous sheath) (Figure 1G). While the majority of these proteins (348) were associated with the nucleus, 59% of the phosphoproteins identified across non-capacitated and capacitated sperm were associated with sperm regions involved in sperm-egg interaction (i.e. the acrosome) and 25% were associated with sperm regions associated with motility (i.e. the axoneme).

### Quantitative phosphopeptide analysis reveals a unidirectional drive in the increase in tyrosine and threonine phosphorylation in human sperm capacitation

Among the 1,773 phosphopeptides identified in this study (Figure 2A), 1,711 were conserved between non-capacitated and capacitated human sperm (96.5%), with an additional 8 phosphopeptides being unique to the non-capacitated and 54 unique to the capacitated samples. These peptides were composed of an average of 2.37 phosphorylated amino acids per peptide (phosphoresidues), with each sperm treatment condition analyzed consisting of over 2,040 phosphorylation events (Figure 2B). Notably, 86 of the observed phosphoproteins had five or more phosphorylated amino acids (Figure 2C), with several examples of hyperphosphorylation including fibrous sheath interacting protein 2 (FSIP2) with 64 phosphoresidues in non-capacitated and 67 phosphoresidues in capacitated sperm (Table 1). Focusing first on the conserved phosphoproteins in both non-capacitated and capacitated sperm, we compared the quantitative differences in phosphopeptide abundance (Figure 2D). This analysis revealed 374 phosphopeptides with significantly altered abundance following sperm capacitation (FC ± 1.5, *p*-value ≤ 0.05; Table S1). Capacitation predominantly resulted in an increased abundance of phosphopeptides (215; 57.5%) with several well characterized motility-associated proteins increasing in phosphorylation (i.e. A kinase anchoring proteins 3 and 4; AKAP3/AKAP4) likely related to the activation of kinases. However, non-capacitated sperm also harbored substantially increased phosphorylation in a subset of proteins (159; 42.5%), potentially related to either the induction or de-repression of phosphatase activity.

**Figure 2:**
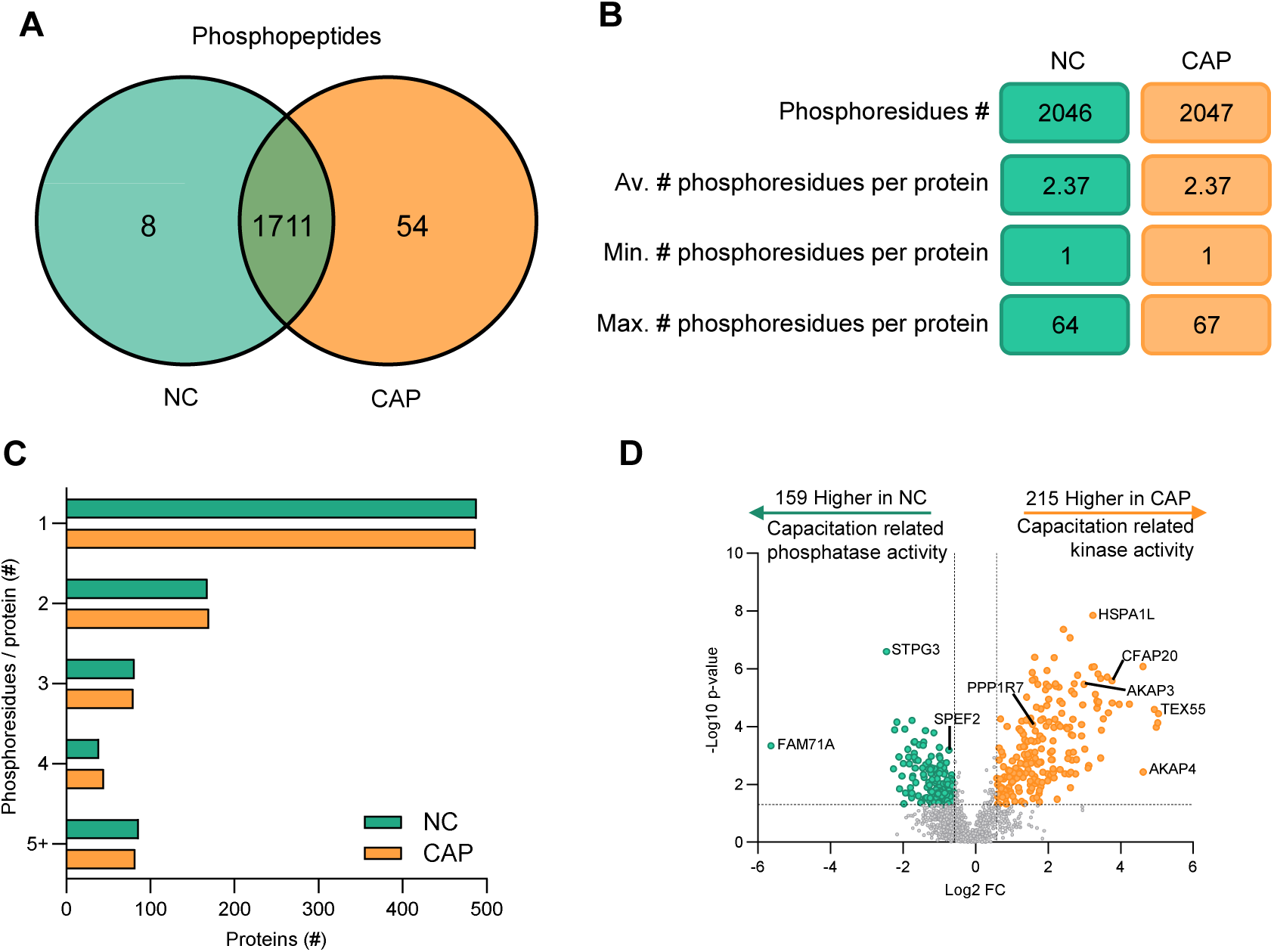
Characterizing the human sperm phosphoproteome; quantitative phosphopeptide analysis. (A) Venn diagram depicting shared and unique phosphopeptides between non-capacitated (NC; green) and capacitated (CAP; orange) populations. (B) The total number of phosphorylated amino acids (phosphoresidues) identified within each populations phosphoproteomic signature, non-capacitated (2,046; green), and capacitated (2,047; orange) along with the average number, minimum number, and maximum number of phosphoresidues per population. (C) Bar graph depicting the number of phosphoresidues/protein. (D) Volcano plot depicting peptides of the conserved phosphoproteome (1711 phosphopeptides) and the differential abundance of 374 phosphopeptides with capacitation; log_2_fold change (x-axis) and – log_10_ *p*-value (y-axis) with thresholds of ± 1.5 fold change and *p*-value ≤ 0.05.

**Table 1:** Top 10 hyper-phosphorylated proteins.

| UniProt | Gene Symbol | Protein name | Phospho-residues (#) |  |  |  |  |  | Uniprot/GO Cellular Localisation |
| --- | --- | --- | --- | --- | --- | --- | --- | --- | --- |
|  |  |  | NC |  |  | CAP |  |  |  |
|  |  |  | pS | pT | pY | pS | pT | pY |  |
| Q5CZC0 | FSIP2 | Fibrous sheath-interacting protein 2 | 60 | 3 | 1 | 60 | 3 | 4 | Midpiece; Axoneme; Flagellum; End Piece; Connecting Piece; Principal Piece |
| Q5JQC9 | AKAP4 | A-kinase anchor protein 4 | 20 | 2 | - | 19 | 2 | - | Nucleus; Midpiece; Flagellum; End Piece; Connecting Piece; Principal Piece; Fibrous Sheath |
| Q8TCU4 | ALMS1 | Alstrom syndrome protein 1 | 23 | - | - | 21 | - | 1 | - |
| Q53TS8-4 | C2CD6 | C2 calcium-dependent domain-containing protein 6 | 23 | - | 1 | 21 | - | 2 | Flagellum |
| Q68DN1 | C2orf16 | Uncharacterized protein C2orf16 | 17 | - | - | 17 | - | - | Nucleus |
| O75969 | AKAP3 | A-kinase anchor protein 3 | 9 | 1 | 1 | 9 | - | 2 | Nucleus; Midpiece; Flagellum; Principal Piece; Fibrous Sheath; Acrosome |
| Q96P26 | NT5C1B | Cytosolic 5'-nucleotidase 1B | 11 | 1 | 1 | 11 | 2 | 2 | Nucleus |
| Q8N5Q1 | FAM71E2 | Protein FAM71E2 | 13 | 1 | - | 12 | 1 | - | - |
| O43236-7 | SEPTIN4 | Isoform 7 of Septin-4 | 7 | - | 1 | 8 | - | 2 | Nucleus; Mitochondria; Annulus; Flagellum; Principal Piece |
| Q8TC71 | SPATA18 | Mitochondria-eating protein | 8 | - | - | 8 | - | - | Mitochondria |
| Total: |  |  | 191 | 8 | 5 | 186 | 8 | 13 |  |

The phosphopeptide with the largest significant fold change captured resides in Protein FAM71A (FAM71A, FC = 50). This peptide was highest in non-capacitated sperm and downregulated during capacitation. However, the subsequent 55 largest significant fold changes were exclusively related to phosphopeptides increased following capacitation; the largest changes among these were PTTG1IP family member 2 (PTTG1IP2; FC = 32.9), Valine-tRNA ligase (VARS1; FC = 32.6), and Cytosolic 5’-nucleotidase 1B (NT5C1B; FC = 31.6). Other notable significantly upregulated phosphopeptides included Testis-specific expressed protein 55 (TEX55; FC = 30.6), A-kinase anchor protein 4 (AKAP4; FC = 24.6), Fibrous sheath-interacting protein 2 (FSIP2; FC = 13.6), Cilia- and flagella-associated protein 20 (CFAP20; FC = 13.6), A-kinase anchor protein 3 (AKAP3; FC = 11.1), and Isoform 7 of Septin-4 (SEPTIN4; FC = 10.5). Following FAM71A, the ensuing most significantly down regulated phosphopeptides (highest in non-capacitated sperm) had notably more modest fold changes, chiefly Protein STPG3 (STPG3; FC 5.5), Eukaryotic translation factor 3 subunit A (EIF3A; FC = 4.8), and Lamin tail domain containing protein 2 (LMNTD2; FC 4.7).

Phosphorylation of amino acids in somatic cells principally occurs in a conserved ratio of 86%:12%:2% for serine, threonine, and tyrosine residues^66^, respectively, as the three most common phosphorylated residues. In non-capacitated human sperm, a ratio of 91.2%:7.0%:1.8% [serine, threonine, and tyrosine] was detected (Figure 3A). In contrast, capacitated human sperm exhibited a 1.7-fold increase in tyrosine phosphorylation with a ratio of 89.8%7.2%:3.0% [serine, threonine, and tyrosine] (Figure 3A). Direct comparison of the phosphopeptides of the two sperm populations identified a subset of conserved peptides with serine (1,302 peptides), threonine (99 peptides), or tyrosine (26 peptides) phosphorylation detected in both non-capacitated and capacitated cells (Figure 3B). A total of 10 phosphoserine peptides and one phosphothreonine were uniquely detected in non-capacitated sperm, indicating the presumptive dephosphorylation of these phosphoresidues during capacitation. Capacitated sperm were represented similarly regarding phosphoserine and phosphothreonine peptides uniquely derived during capacitation, with 25 phosphoserine and 7 phosphothreonine peptides detected only in capacitated sperm cells. Interestingly, no phosphotyrosine events were detected exclusive to non-capacitated sperm. Conversely, 18 phosphopeptides with tyrosine phosphorylation were exclusively detected following capacitation of human sperm (Figure 3B and Table 2) demonstrating a unidirectional shift in tyrosine phosphorylation signaling in human sperm.

**Figure 3:**
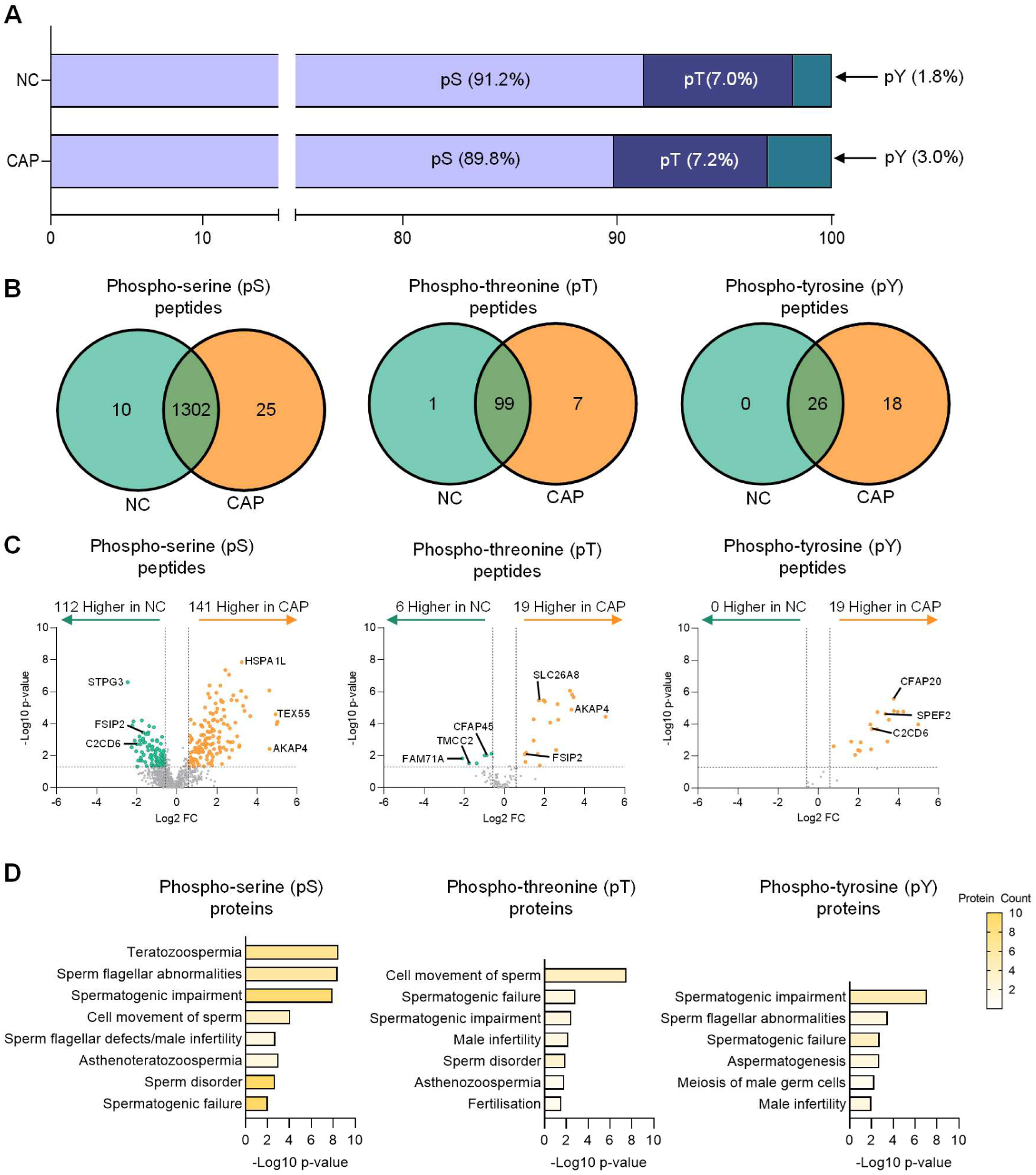
Characterizing human sperm phosphoproteome: phosphoresidue distributions. (A) Bar graph depicting the distribution of Serine (pS), Threonine (pT), and Tyrosine (pY) phosphorylation between non-capacitated (NC) and capacitated (CAP) populations.(B) Venn diagrams depicting shared and unique phosphopeptides between non-capacitated (NC; green) and capacitated (CAP; orange) sperm populations for phosphoserine (pS), phosphothreonine (pT), and phosphotyrosine (pY). (C) Volcano plot depicting peptides of the conserved phosphopeptides for phosphoserine (pS; 1302), phosphothreonine (pT; 99), and phosphotyrosine (pY; 26); log_2_ fold change (x-axis) and –log10 *p*-value (y-axis) with thresholds of ±1.5 fold change and *p*-value ≤ 0.05. (D) Most highly enriched disease and functions identified by Ingenuity Pathway Analysis (IPA); enrichment determined by *p* ≤ 0.05, -log_10_ *p*-value (x-axis).

**Table 2:**
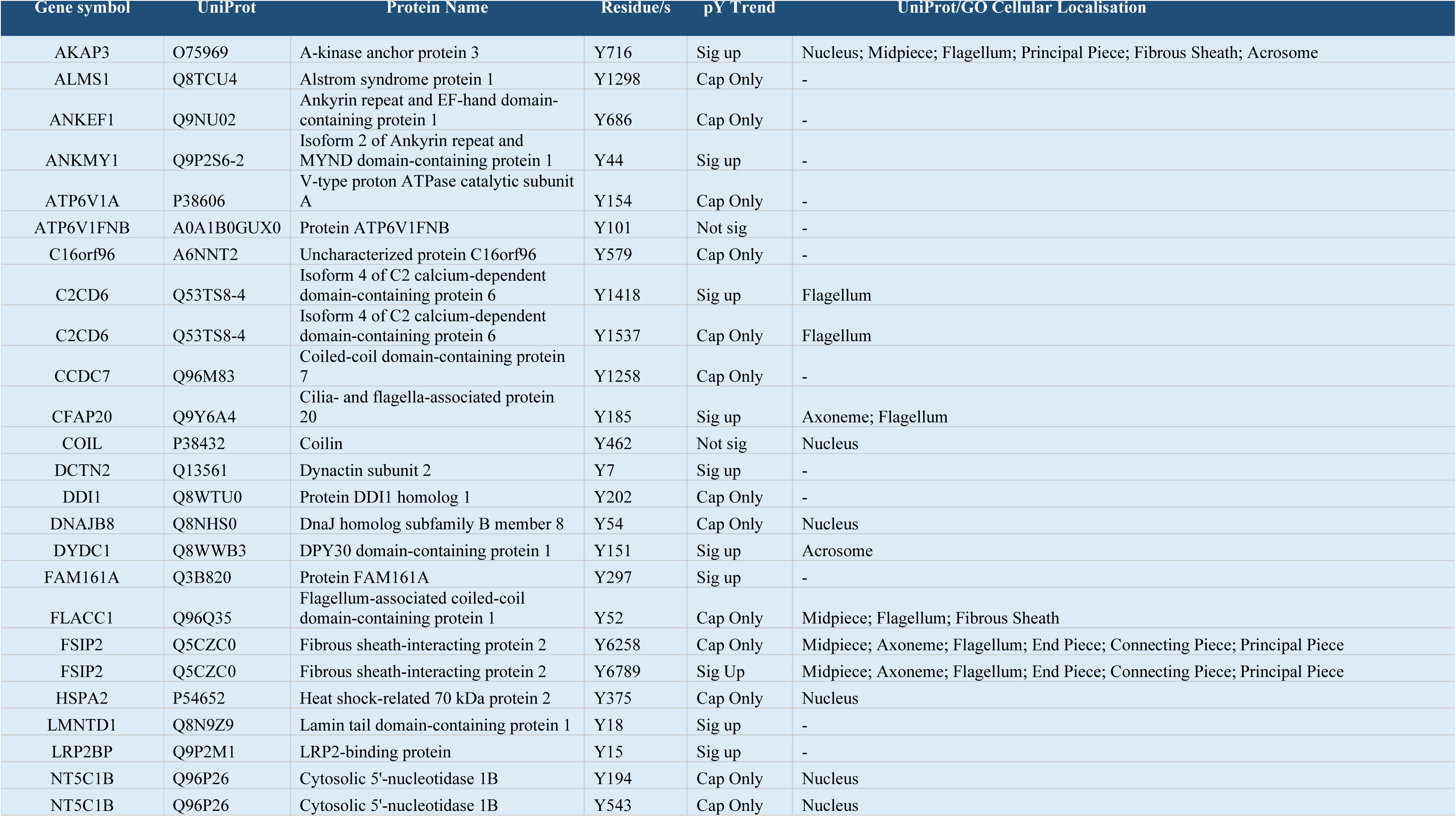
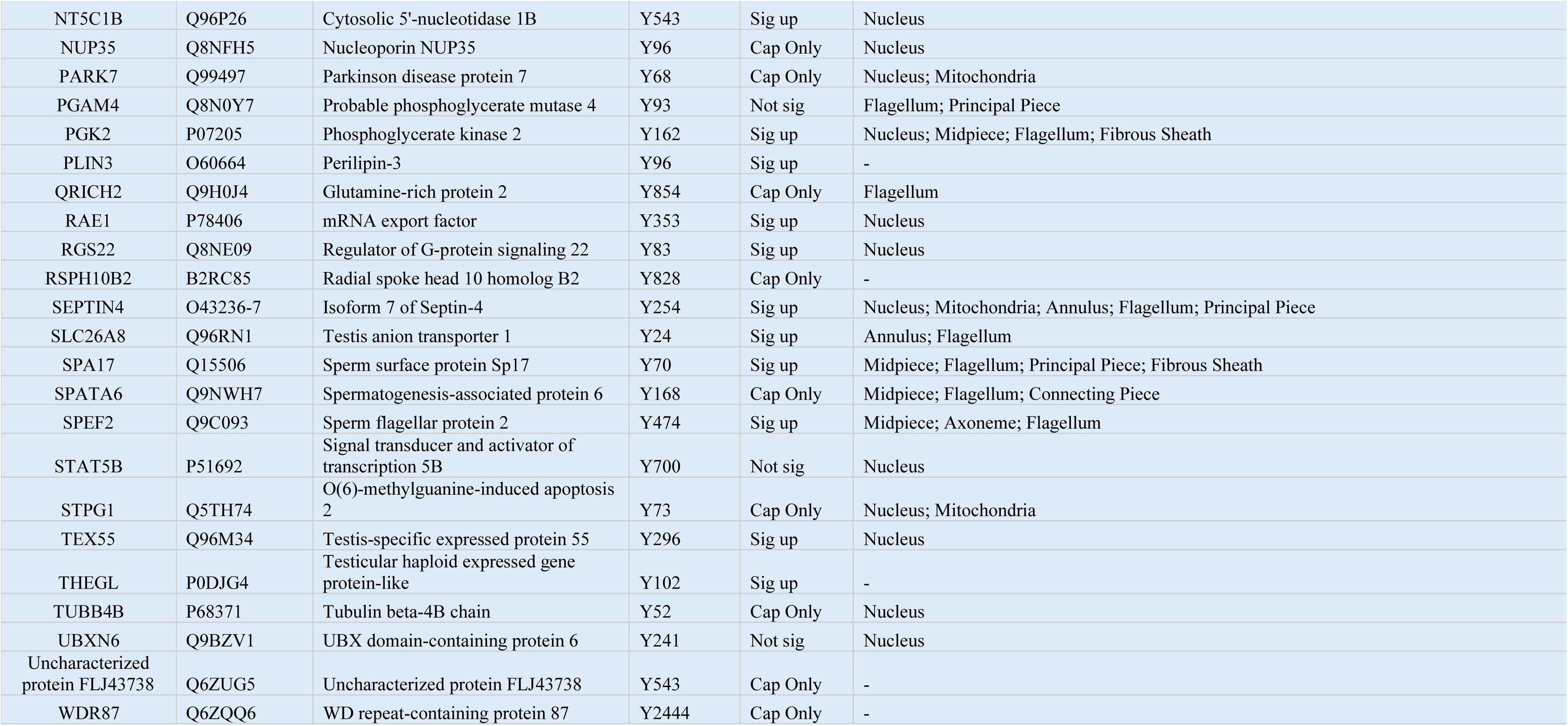
Tyrosine phosphorylated proteins in non-capacitated and capacitated human sperm; their phospho-sites, and phospho-status relative to capacitation. “Cap Only” represents detection in only capacitated samples, “Sig Up” represents significantly upregulated in capacitated samples, and “Not Sig” represents no capacitation dependent change detected.

| Gene symbol | UniProt | Protein Name | Residue/s | pY Trend | UniProt/GO Cellular Localisation |
| --- | --- | --- | --- | --- | --- |
| AKAP3 | O75969 | A-kinase anchor protein 3 | Y716 | Sig up | Nucleus; Midpiece; Flagellum; Principal Piece; Fibrous Sheath; Acrosome |
| ALMS1 | Q8TCU4 | Alstrom syndrome protein 1 | Y1298 | Cap Only | - |
| ANKEF1 | Q9NU02 | Ankyrin repeat and EF-hand domain-containing protein 1 | Y686 | Cap Only | - |
| ANKMY1 | Q9P2S6-2 | Isoform 2 of Ankyrin repeat and MYND domain-containing protein 1 | Y44 | Sig up | - |
| ATP6V1A | P38606 | V-type proton ATPase catalytic subunit A | Y154 | Cap Only | - |
| ATP6V1FNB | A0A1B0GUX0 | Protein ATP6V1FNB | Y101 | Not sig | - |
| C16orf96 | A6NNT2 | Uncharacterized protein C16orf96 | Y579 | Cap Only | - |
| C2CD6 | Q53TS8-4 | Isoform 4 of C2 calcium-dependent domain-containing protein 6 | Y1418 | Sig up | Flagellum |
| C2CD6 | Q53TS8-4 | Isoform 4 of C2 calcium-dependent domain-containing protein 6 | Y1537 | Cap Only | Flagellum |
| CCDC7 | Q96M83 | Coiled-coil domain-containing protein 7 | Y1258 | Cap Only | - |
| CFAP20 | Q9Y6A4 | Cilia- and flagella-associated protein 20 | Y185 | Sig up | Axoneme; Flagellum |
| COIL | P38432 | Coilin | Y462 | Not sig | Nucleus |
| DCTN2 | Q13561 | Dynactin subunit 2 | Y7 | Sig up | - |
| DDI1 | Q8WTU0 | Protein DDI1 homolog 1 | Y202 | Cap Only | - |
| DNAJB8 | Q8NHS0 | DnaJ homolog subfamily B member 8 | Y54 | Cap Only | Nucleus |
| DYDC1 | Q8WWB3 | DPY30 domain-containing protein 1 | Y151 | Sig up | Acrosome |
| FAM161A | Q3B820 | Protein FAM161A | Y297 | Sig up | - |
| FLACC1 | Q96Q35 | Flagellum-associated coiled-coil domain-containing protein 1 | Y52 | Cap Only | Midpiece; Flagellum; Fibrous Sheath |
| FSIP2 | Q5CZC0 | Fibrous sheath-interacting protein 2 | Y6258 | Cap Only | Midpiece; Axoneme; Flagellum; End Piece; Connecting Piece; Principal Piece |
| FSIP2 | Q5CZC0 | Fibrous sheath-interacting protein 2 | Y6789 | Sig Up | Midpiece; Axoneme; Flagellum; End Piece; Connecting Piece; Principal Piece |
| HSPA2 | P54652 | Heat shock-related 70 kDa protein 2 | Y375 | Cap Only | Nucleus |
| LMNTD1 | Q8N9Z9 | Lamin tail domain-containing protein 1 | Y18 | Sig up | - |
| LRP2BP | Q9P2M1 | LRP2-binding protein | Y15 | Sig up | - |
| NT5C1B | Q96P26 | Cytosolic 5'-nucleotidase 1B | Y194 | Cap Only | Nucleus |
| NT5C1B | Q96P26 | Cytosolic 5'-nucleotidase 1B | Y543 | Cap Only | Nucleus |
| NT5C1B | Q96P26 | Cytosolic 5'-nucleotidase 1B | Y543 | Sig up | Nucleus |
| NUP35 | Q8NFH5 | Nucleoporin NUP35 | Y96 | Cap Only | Nucleus |
| PARK7 | Q99497 | Parkinson disease protein 7 | Y68 | Cap Only | Nucleus; Mitochondria |
| PGAM4 | Q8N0Y7 | Probable phosphoglycerate mutase 4 | Y93 | Not sig | Flagellum; Principal Piece |
| PGK2 | P07205 | Phosphoglycerate kinase 2 | Y162 | Sig up | Nucleus; Midpiece; Flagellum; Fibrous Sheath |
| PLIN3 | O60664 | Perilipin-3 | Y96 | Sig up | - |
| QRICH2 | Q9H0J4 | Glutamine-rich protein 2 | Y854 | Cap Only | Flagellum |
| RAE1 | P78406 | mRNA export factor | Y353 | Sig up | Nucleus |
| RGS22 | Q8NE09 | Regulator of G-protein signaling 22 | Y83 | Sig up | Nucleus |
| RSPH10B2 | B2RC85 | Radial spoke head 10 homolog B2 | Y828 | Cap Only | - |
| SEPTIN4 | O43236-7 | Isoform 7 of Septin-4 | Y254 | Sig up | Nucleus; Mitochondria; Annulus; Flagellum; Principal Piece |
| SLC26A8 | Q96RN1 | Testis anion transporter 1 | Y24 | Sig up | Annulus; Flagellum |
| SPA17 | Q15506 | Sperm surface protein Sp17 | Y70 | Sig up | Midpiece; Flagellum; Principal Piece; Fibrous Sheath |
| SPATA6 | Q9NWH7 | Spermatogenesis-associated protein 6 | Y168 | Cap Only | Midpiece; Flagellum; Connecting Piece |
| SPEF2 | Q9C093 | Sperm flagellar protein 2 | Y474 | Sig up | Midpiece; Axoneme; Flagellum |
| STAT5B | P51692 | Signal transducer and activator of transcription 5B | Y700 | Not sig | Nucleus |
| STPG1 | Q5TH74 | O(6)-methylguanine-induced apoptosis 2 | Y73 | Cap Only | Nucleus; Mitochondria |
| TEX55 | Q96M34 | Testis-specific expressed protein 55 | Y296 | Sig up | Nucleus |
| THEGL | P0DJG4 | Testicular haploid expressed gene protein-like | Y102 | Sig up | - |
| TUBB4B | P68371 | Tubulin beta-4B chain | Y52 | Cap Only | Nucleus |
| UBXN6 | Q9BZV1 | UBX domain-containing protein 6 | Y241 | Not sig | Nucleus |
| Uncharacterized protein FLJ43738 | Q6ZUG5 | Uncharacterized protein FLJ43738 | Y543 | Cap Only | - |
| WDR87 | Q6ZQQ6 | WD repeat-containing protein 87 | Y2444 | Cap Only | - |

Comparing the quantitative differences in phosphoresidue abundance across the three phosphoresidues (Figure 3C and Table S1), analysis of phosphoserine revealed 253 / 1302 (19.4%) phosphopeptides significantly altered by capacitation (FC ± 1.5, *p*-value ≤ 0.05). Among the dysregulated phosphoserine phosphopeptides, 141 were significantly increased following capacitation, including 45 of the 50 peptides with the greatest fold changes observed. Proteins with increased phosphoserine phosphopeptides included Putative protein FAM10A4 (ST13P4; FC = 24.5) and Coiled-coil domain-containing protein 63 (CCDC63; FC = 12.7), as well as previously mentioned VARS1 (FC = 32.6), NT5C1B (FC = 31.6), TEX55 (FC = 30.6), and AKAP4 (FC = 24.6). In non-capacitated sperm, 112 phosphoserine peptides were significantly higher [prior to capacitation], the top five of which mapped to Protein STPG3 (STPG3; FC = 5.5), Eukaryotic translation initiation factor 3 subunit A (EIF3A; FC = 4.8), Lamin tail domain-containing protein 2 (LMNTD2; FC = 4.7), EF-hand calcium -binding domain-containing protein 5 (EFCAB5; FC = 4.5), and serine/threonine-protein phosphatase 4 regulatory subunit 1 (PPP4R1; FC = 4.3).

Analysis of the conserved phosphothreonine peptides in non-capacitated and capacitated human sperm identified 25 / 99 (25.3%) significantly altered peptides (Figure 3C, Table S1). Capacitation increased 19 phosphothreonine peptides from proteins including PTTG1P family member 2 (PTTG1IP2; FC = 32.9), Endoplasmic reticulum-Golgi intermediate compartment protein 2 (ERGIC2; FC = 11), dynein axonal intermediate chain 1 (DNAI1; FC = 10.4), AKAP4 (FC = 10.2), and Glutamine synthetase (GLUL; FC = 9.6). Phosphothreonine peptides were underrepresented in non-capacitated sperm, with six threonine phosphopeptides significantly higher prior to capacitation, mapping to proteins including FAM71A (FC = 4.3), Transmembrane and coiled-coil domains protein 2 (TMCC2; FC = 3.4), protein flattop (CFAP126; FC = 2.6), and Cilia- and flagella-associated protein 45 (CFAP45; FC = 1.9). Analysis of phosphotyrosine peptides revealed 19 / 28 (67.9%) significantly altered peptides, and uniquely, all 19 were higher as a consequence of capacitation. Phosphotyrosine peptides were derived from proteins including DPY30 domain-containing protein 1 (PYDC1; FC = 18.9), Cilia- and flagella-associated protein 20 (CFAP20; FC = 13.6), as well as previously mentioned NT5C1B (FC = 31.6), TEX55 (FC = 15.6), and FSIP2 (FC = 13.6). The distribution of all phospho-modifications is presented in Figure S2A.

To better appreciate the biological processes regulated by phosphorylation in human sperm, we further interrogated the subset of phosphoproteins that were either significantly altered (down-and up-regulated) and those exclusive to either non-capacitated or capacitated sperm using Ingenuity Pathway Analysis (IPA) (Table S2). Focusing on reproductive and sperm-related functions (Figure 3D), we observed significant enrichment (*p* ≤ 0.05) in the phosphorylation of proteins implicated in spermatogenesis, featuring testicular derived pathologies including spermatogenic failure (phosphoserine, phosphothreonine, and phosphotyrosine), spermatogenic impairment (phosphoserine, phosphothreonine, phosphotyrosine), teratozoospermia (phosphoserine), meiosis of germ cells (phosphotyrosine), and aspermatogenesis (phosphotyrosine). Significant enrichment (*p* ≤ 0.05) of phosphorylated proteins integral to the sperm flagellum and motility were also prominent with motility associated pathologies and sperm flagellar abnormalities (phosphoserine and phosphotyrosine), cell movement of sperm (phosphoserine and phosphothreonine), and asthenozoospermia (phosphothreonine). Taking into consideration the unidirectional nature of the tyrosine phosphorylation detected and that tyrosine phosphorylation is a well cited hallmark of capacitation ^37,38,63–65^, Table 2 features a curated list of tyrosine modified peptides, the proteins they constitute, their gene ontology and notation of those proteins with previously characterized roles in male fertility.

### In-silico analysis of human sperm phosphoproteins reveals putative kinase candidates with potential roles in human sperm capacitation

To contextualize the human sperm phosphoproteome in relation to known phosphosites identified in all biological systems, we used PhosphoSitePlus; a comprehensive repository for protein post-translational modifications, in particular protein phosphorylation. To conduct this comparison, a refinement of the phosphoproteome was necessary to include only peptides where the exact site of phosphorylation was identifiable, yielding 669 phosphoproteins with a total of 1,230 phosphoresidues. A direct comparison of these phosphoproteins to the PhosphoSitePlus repository ^67,68^ revealed that of the 1,230 phosphoresidues, 755 sperm phosphoresidues (63%) were reported for the first time in any cell type (Figure 4A) with an additional 49 sperm phosphoproteins that had not been previously identified as phosphorylated in other studies (Figure S2B).

**Figure 4:**
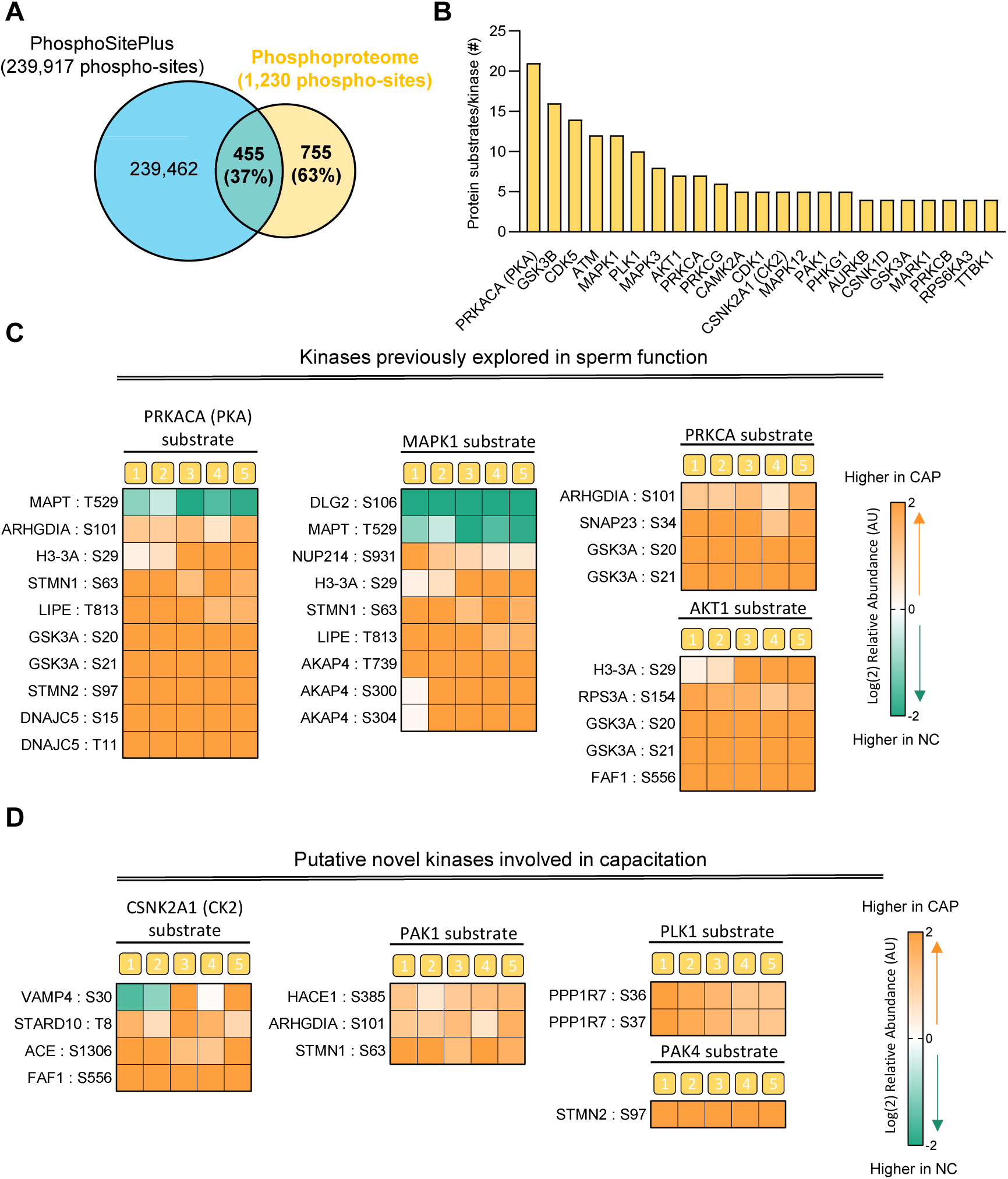
In-silico analysis of human sperm phosphoproteins and kinase prediction using Phosphomatics. (A) Venn diagram of shared and unique phosphoresidues (1230 identified amino acids/sites) compared to PhosphoSite Plus database (239,917 sites), 755 (63%) phosphoresidues not previously reported (PhosphoSite Plus accessed 11/09/2023). (B) Bar graph of kinases with ≥4 protein substrates detected in at least one population (non-capacitated and/or capacitated sperm; PhosphoSitePlus accessed 02/03/2023). (C) Heat map of previously characterized kinases in sperm and their substrates with significantly upregulated phosphorylation; higher in non-capacitated (green) or capacitated (orange) human sperm samples, constructed from Phosphomatics (accessed 05/04/2023). (D) Heat map of previously not reported candidate kinases in sperm and their substrates with significantly upregulated phosphorylation; higher in non-capacitated (green) or capacitated (orange) human sperm samples, constructed from Phosphomatics (accessed 05/04/2023).

We then explored these data further at the phosphoprotein level to shed light on the kinases that may have roles in sperm development. Employing RoKAI, a computational tool for inferring kinase activity using functional networks, we observed 18 kinases with ≥ 2 protein substrates within the sperm phosphoproteome (Figure S2C), including cAMP-dependent protein kinase catalytic subunit alpha (PRKACA; substrates = 12), Mitogen-activated protein kinase 1 (MAPK1; substrates = 6), and Serine/threonine protein kinase 1 (PAK1; substrates = 5). Complementing this analysis, the use of an alternative in silico phosphoproteomic data analysis tool, Phosphomatics, also identified 23 kinases with ≥ 4 protein substrates within the human sperm phosphoproteome (Figure 4B), including PRKACA (substrates = 21), Glycogen synthase kinase-3 beta (GSK3B; substrates = 16), Cyclin-dependent kinase 5 (CDK5; substrates = 14).

Notwithstanding the large portion of the sperm phosphoresidues that reside in previously undescribed locations, we further interrogated the phosphoresidue data using RoKAI and Phosphomatics to derive biological insight into which kinases may be active under capacitation conditions. Phosphomatics enabled motif analysis of the amino acids located ±7 residues from the phosphoresidue in each peptide (Figure S2D). From this analysis, the dominant motif observed was that of PKA, a finding that resonates with this kinase being the most well characterized regulator of sperm capacitation. Expanding this analysis, RoKAI kinase activity inference predicted the activity of 10 kinases (Figure S2E), key among them being Inhibitor of nuclease factor kappa-B kinase subunit epsilon (IKBKE) in terms of both activity score (5.670) and significance evaluation (Z-score = 5.124). PKA, cAMP-dependent protein kinase catalytic subunit alpha (PRKACA or PKA C-alpha) also featured, with a kinase-activity score of 2.42 (Z-score = 2.187). Further, Phosphomatics kinase inference identified 48 kinases with 31 significantly altered phosphoresidue substrates between the non-capacitated and capacitated human sperm samples (Table S3). Notwithstanding the promiscuity of several phosphoresidues, which are capable of being phosphorylated by numerous kinases, we further mined these kinases and their substrates for previous associations to sperm function and fertility using UniProt. Among those with previously identified roles in fertility (Figure 4C), PRKACA was the highest in terms of number of substrates, with 9 significantly upregulated phosphoresidues mapping to 7 protein substrates following capacitation. Other kinases with known roles in capacitation also had several phosphoresidues significantly upregulated following capacitation (Figure 4C), including MAPK1 (7 residues, 4 proteins), Protein kinase C alpha type (PRKCA; 4 residues, 3 proteins), and AKT1 (5 residues, 4 proteins).

Importantly, several kinase-substrate relationships were revealed through this approach where the kinases mapped had no previous known role in sperm function. Specifically, these kinases were CK2 (4 residues, 4 proteins), PAK1 (3 residues, 3 proteins), PLK1 (2 residues, 1 protein), and PAK4 (1 residue, 1 protein) (Figure 4D). Superimposed onto a human kinome map (Figure 5) is a visual representation of all the kinases identified that had previous associations to sperm function (41 kinases), the kinases detected within our phosphoproteomic profiling of human sperm (20 kinases), and the kinases predicted using our *in-silico* analysis strategy (52 kinases) [for a further annotated kinase map, see Figure S3].

**Figure 5:**
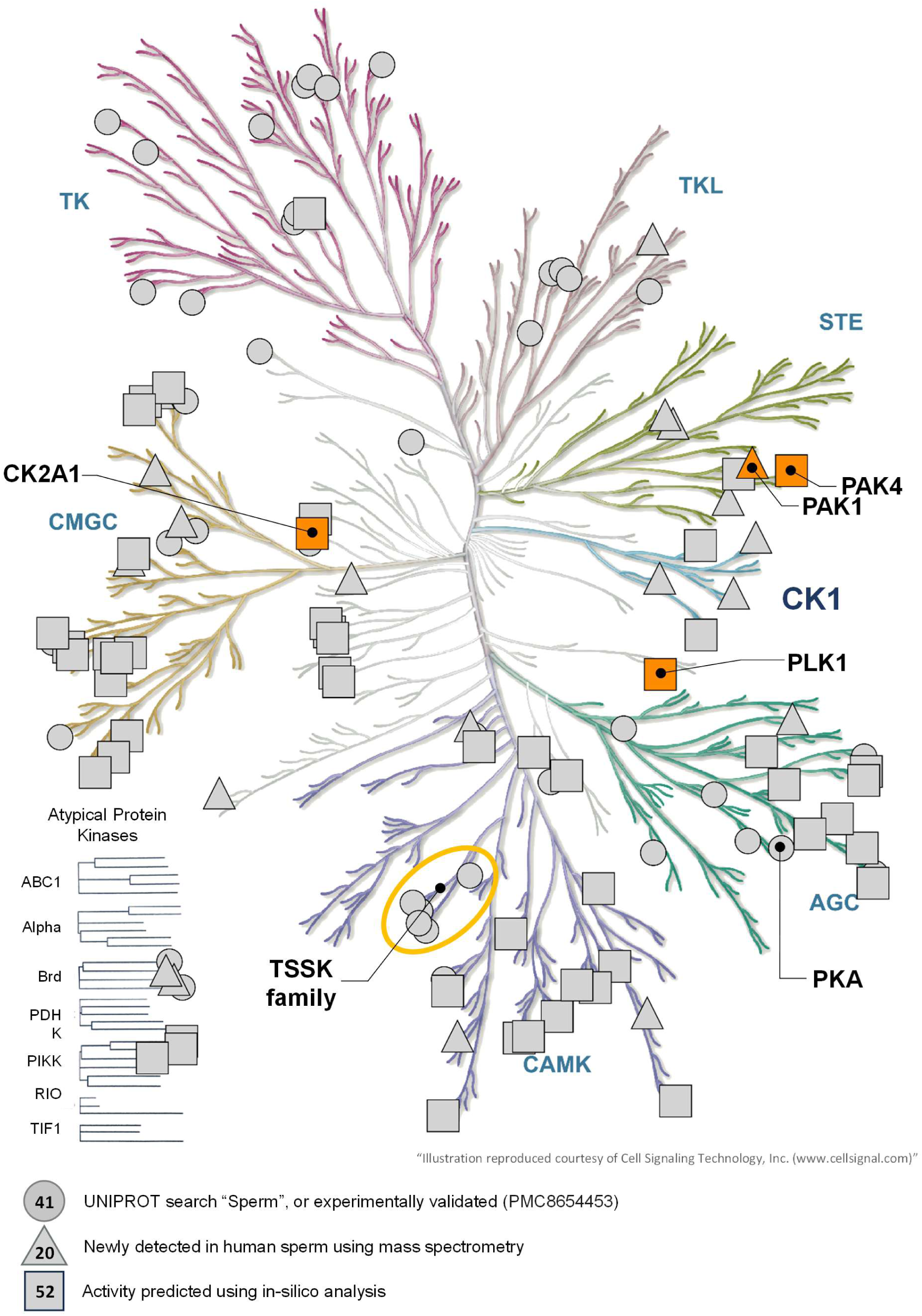
The human sperm kinome. “Sperm” associated kinases from UniProt and experimentally validated kinases (Circle), kinases detected using mass-spectrometry (this study; Triangle), and kinases predicted using Phosphomatics *in-silico* analysis (this study; Square) represented on the human kinome (Cell Signaling Technology, Inc.). Shown in orange are kinases inhibited in this study. Where a kinase occurred in two or more groups it was assigned to the highest level of evidence where previous validation (UniProt) < mass spectrometry < in-silico predicted. CAMK; Calcium/calmodulin-dependent protein kinase, AGC; Containing PKA, PKG, PKC families, CK1; Casein kinase 1 family, STE; Homologs of yeast Sterile 7, Sterile 11, Sterile 20 kinases, TKL; Tyrosine kinase-like, TK; Tyrosine kinase family, CMGC; Containing CDK, MAPK, GSK3, CLK families.

Based on these combined analyses, a subset of kinases with no previous functional data relating to sperm function were selected for targeted pharmacological inhibition to directly investigate their contributions to human sperm capacitation (Figure 4D). More specifically, CK2 was selected in reference to it being the only kinase reported to phosphorylate Angiotensin-converting enzyme (ACE), a protein with a testis-specific isoform, expressed in spermatocytes in the adult testis, and required for male fertility including reported roles in sperm-oocyte interaction ^69^. PAK1, serine/threonine-protein kinase PAK2 (PAK2) and PAK4 all had significantly altered substrates in our dataset, and PAK1 is among the kinase candidates that phosphorylates Rho GDP-dissociation inhibitor 1 (ARHGDIA), previously reported to be decreased in the sperm of IVF patients with low fertilization rates, and significantly increased in the sperm of normozoospermic men and IVF patients with higher fertilization rates ^70^. Finally, PLK1 was the only kinase known to phosphorylate protein phosphatase 1 regulatory subunit 7 (PPP1R7), a regulatory subunit of serine/threonine-protein phosphatase PP1 (PPP1A) ^71,72^. PPP1A is a phosphatase with a testis-specific isoform (PP1γ2) enriched in sperm during epididymal maturation, with de-capacitation like activity ^72^. The inhibition of PP1γ2 by PPP1R7, and other regulatory subunits, is phosphorylation dependent, and as such we hypothesized that PLK1 may play an unappreciated role in sperm capacitation ^71^.

### In vitro kinase inhibition during human sperm capacitation reveals a functional role for PLK1 in regulating capacitation dependent protein tyrosine phosphorylation

To confirm whether the identified kinases participate in sperm capacitation, kinases were targeted through pharmacological inhibition over a 3.5 hour time course. Beginning with a 30 min pre-incubation in non-capacitating BWW with or without inhibitors, sperm cells were then centrifuged and resuspended in capacitating BWW ± inhibitors and subjected to capacitating conditions for three hours. Four kinases were targeted by selective inhibitors as follows: CK2 inhibited by CX-4945, PAK1 inhibited by NVS-PAK1-1, PAK4 inhibited by GNE 2861, and PLK1 inhibited by MLN0905. Human sperm cell viability was not perturbed in non-capacitating BWW for 30 min (Figure S4A) or capacitating BWW for 3 hours (Figure 6A) by any of the inhibitors at concentrations from 0.2 μM to 50 μM. Sperm cell motility kinematics were assessed using computer assisted sperm analysis (CASA), and similarly, we demonstrated that kinase inhibition had no overt effect on total motility in either non-capacitating or capacitating conditions over 3.5 hours for all inhibitors assayed (Figure S4B and Figure 6B). Under non-capacitating conditions (30 min) progressive motility was not perturbed by CK2, PAK1 or PLK1 inhibition (Figure S4C). Under the same conditions, inhibition of PAK4 with GNE 2861 at 7.5 μM and 50 μM significantly increased the progressive motility of human sperm (Figure S4C). Under capacitating conditions (Figure 6C), progressive motility significantly increased, as expected, compared to the non-capacitated control (3 hours; Figure 6C). Progressive motility was significantly decreased under the same capacitating conditions in the presence of 50 μM NVS-PAK1-1, 50 μM GNE 2861, and 20 μM MLN0905, inhibiting PAK1, PAK4, and PLK1 respectively (Figure 6C). Notably, these observed decreases in progressive motility fall to levels congruent with non-capacitated sperm. In contrast, CX-4945 (1 μM–50 μM) directed inhibition of CK2 did not affect progressive motility under capacitating conditions. Upon exploring sperm kinematics under capacitating conditions using CASA, several motility parameters were significantly reduced to or below the level of non-capacitated sperm in the presence of PAK1, PAK4, and PLK1 inhibitors, namely straight-line velocity (VSL; Figure S5A), curvilinear velocity (VCL; Figure S5B), and linearity (LIN; Figure S5C), while beat cross frequency (BCF; S5D) increased under conditions of PAK1 inhibition, though remained unperturbed by PAK4 and PLK1 inhibition.

**Figure 6:**
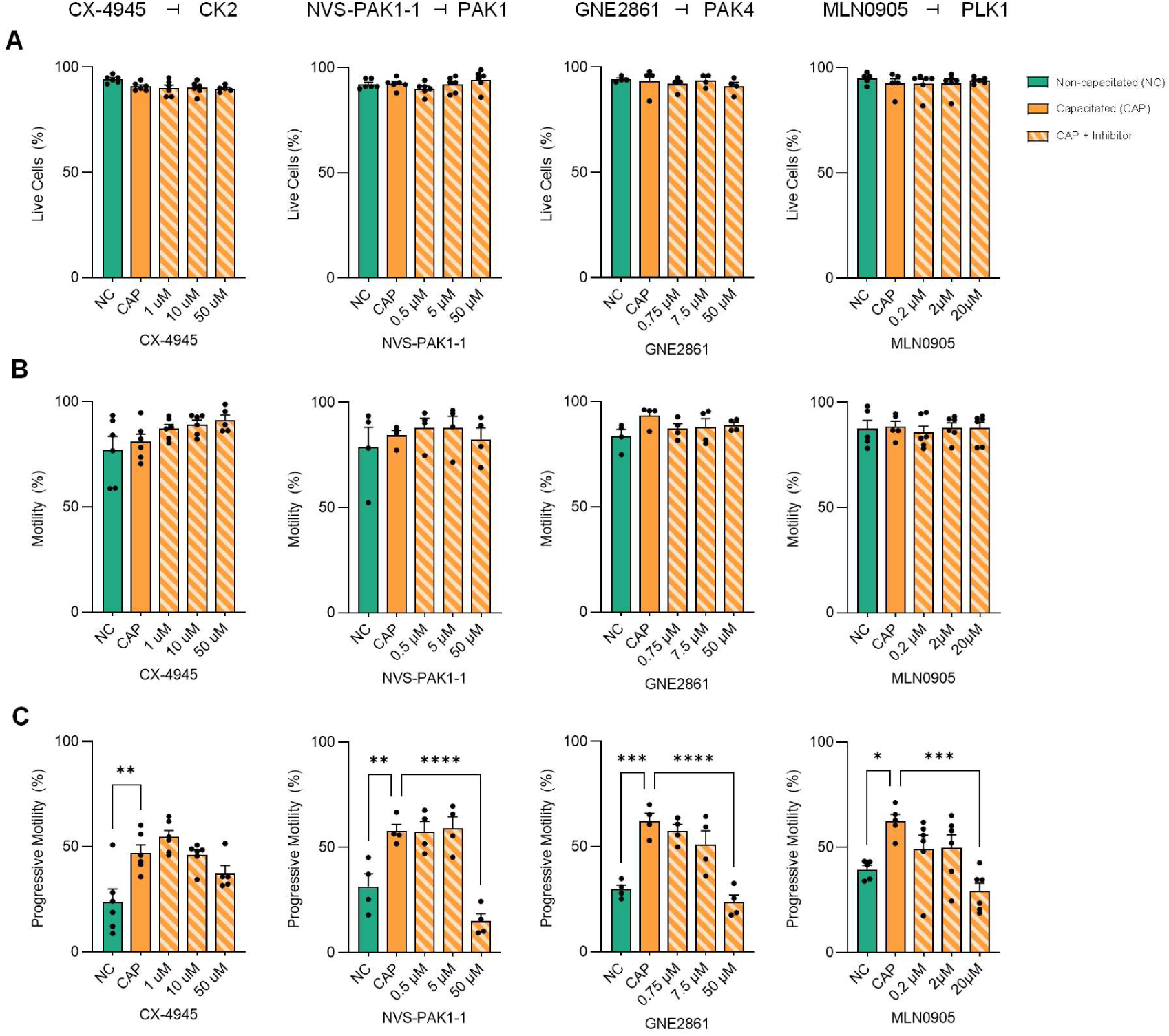
Assessment of the effect of *in vitro* kinase inhibition on sperm viability and motility parameters. Sperm suspensions in non-capacitating BWW were “spiked” with kinase inhibitors to relevant concentrations and pre-incubated for 30 min before non-capacitated (NC; green) results were recorded. Samples were subsequently centrifuged and resuspended in capacitating BWW with relevant DMSO vehicle (CAP; orange) or kinase inhibitor concentrations (orange striped), incubated for 3 hours (37°C, 5% CO_2_), and subjected to (A) viability (eosin) and (B-C) motility assessment (CASA). Data were subject to a non-parametric one-way ANOVA, with Dunnett’s multiple comparisons test; * *p* ≤ 0.05, ** *p* ≤ 0.01, *** *p* ≤ 0.001, **** *p* ≤ 0.0001. All graphs contain n = 4-6 biological samples.

To investigate whether human sperm were still capable of undergoing capacitation in the presence of the four kinase inhibitors, SDS-PAGE and immunoblotting for protein tyrosine phosphorylation was performed. CK2 and PAK1 inhibition (10 μM CX-4945 and 5 μM NVS-PAK1-1 respectively) did not moderate the ability of human sperm to undergo capacitation whereas tyrosine phosphorylation was modestly decreased in sperm with PAK4 inhibition (7.5 μM GNE 2861; Figure 7A and B and Figure S6A and B). Strikingly, PLK1 inhibition (20 μM MLN0905) appeared to prevent the capacitation of human sperm, with tyrosine phosphorylation reduced to a level equivalent to that detected in non-capacitated sperm cells (Figure 7A and B). To further validate this result, human sperm were treated with MLN0905 (PLK1 inhibitor) during capacitation and isolated sperm cells were fixed and probed with an anti-phosphotyrosine antibody (Figure 8A). This experiment confirmed that phosphotyrosine sperm tail labeling was significantly reduced (3.5fold) through PLK1 inhibition (Figure 8B), indicating that PLK1 plays a role in human sperm capacitation signaling. Neither phosphoserine nor phosphothreonine immunodetection was visibly altered in the presence of the kinase inhibitors investigated here (Figure S7A and B, Figure S8A and B). To further assess the functional role of PLK1 in human sperm, we evaluated the ability for human sperm to undergo the acrosome reaction (a critical step preceding fertilization) following PLK1 inhibition. Notably, the inhibition of PLK1 proved to significantly suppress both the spontaneous and progesterone induced acrosome reaction rates, retaining acrosome reaction levels equivalent to non-capacitated sperm in both instances (Figure 8C and Figure S9). With this functional evidence as the impetus, immunolabelled mature human sperm show a midpiece and equatorial localization of PLK1 (Figure S10). This was reinforced by co-labelling with principle piece (α-Tubulin and AKAP3) and midpiece (Glycerokinase) antibodies.

**Figure 7:**
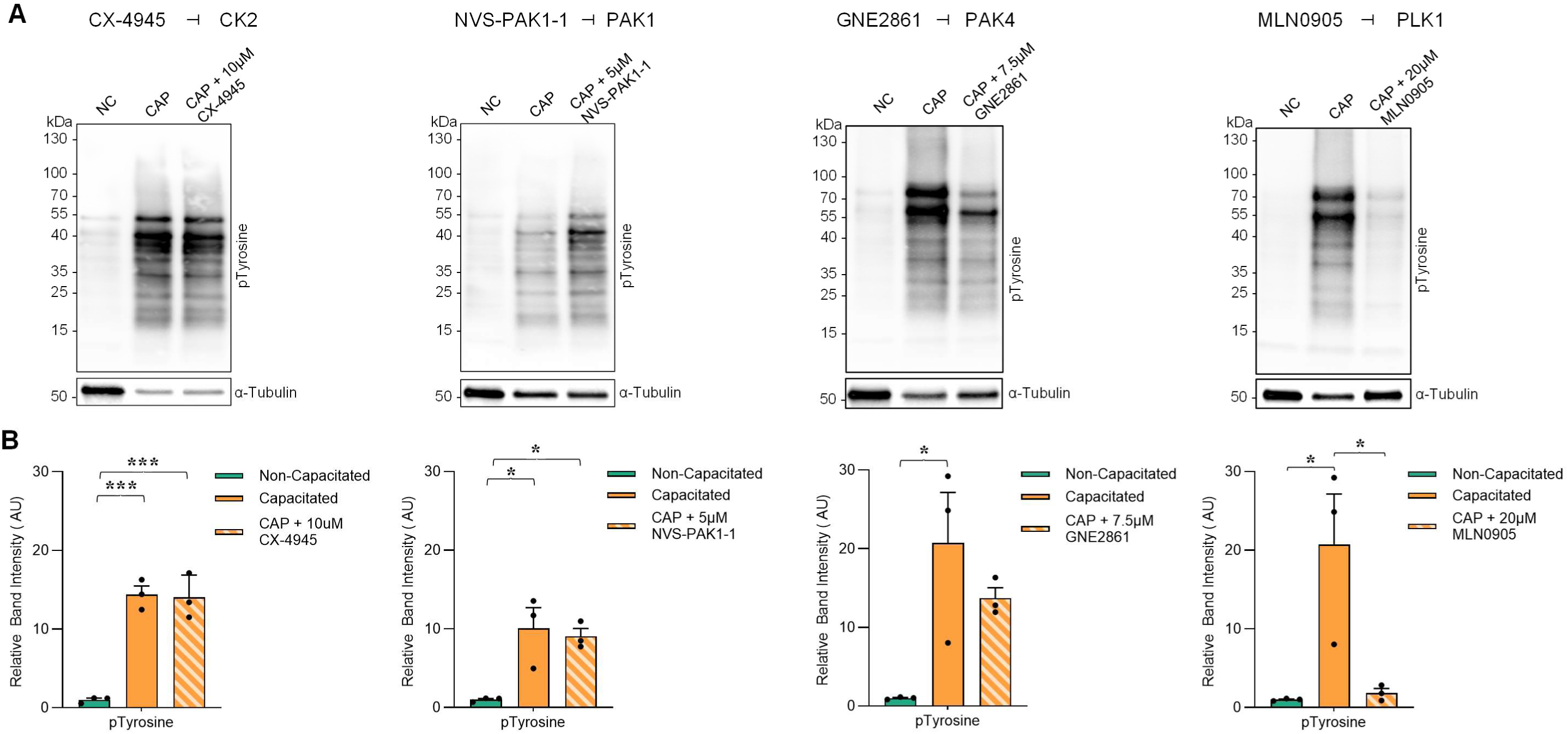
Assessment of the effect of *in vitro* kinase inhibition on sperm capacitation ability. Sperm were incubated (3 hours 37°C, 5% CO_2_) in capacitating BWW with relevant DMSO vehicle (CAP; orange) or kinase inhibitor concentrations (orange striped), and snap frozen along-side non-capacitated sperm (NC; green). To assess capacitation status sperm samples were subjected to (A) SDS-PAGE and immunoblotting for anti-phosphotyrosine (PT66) and subsequent densitometric assessment (B). Data were subject to one-way ANOVA with Tukey’s multiple comparisons test; * *p* ≤ 0.05, *** *p* ≤ 0.001. All graphs contain n = 3 biological samples.

**Figure 8:**
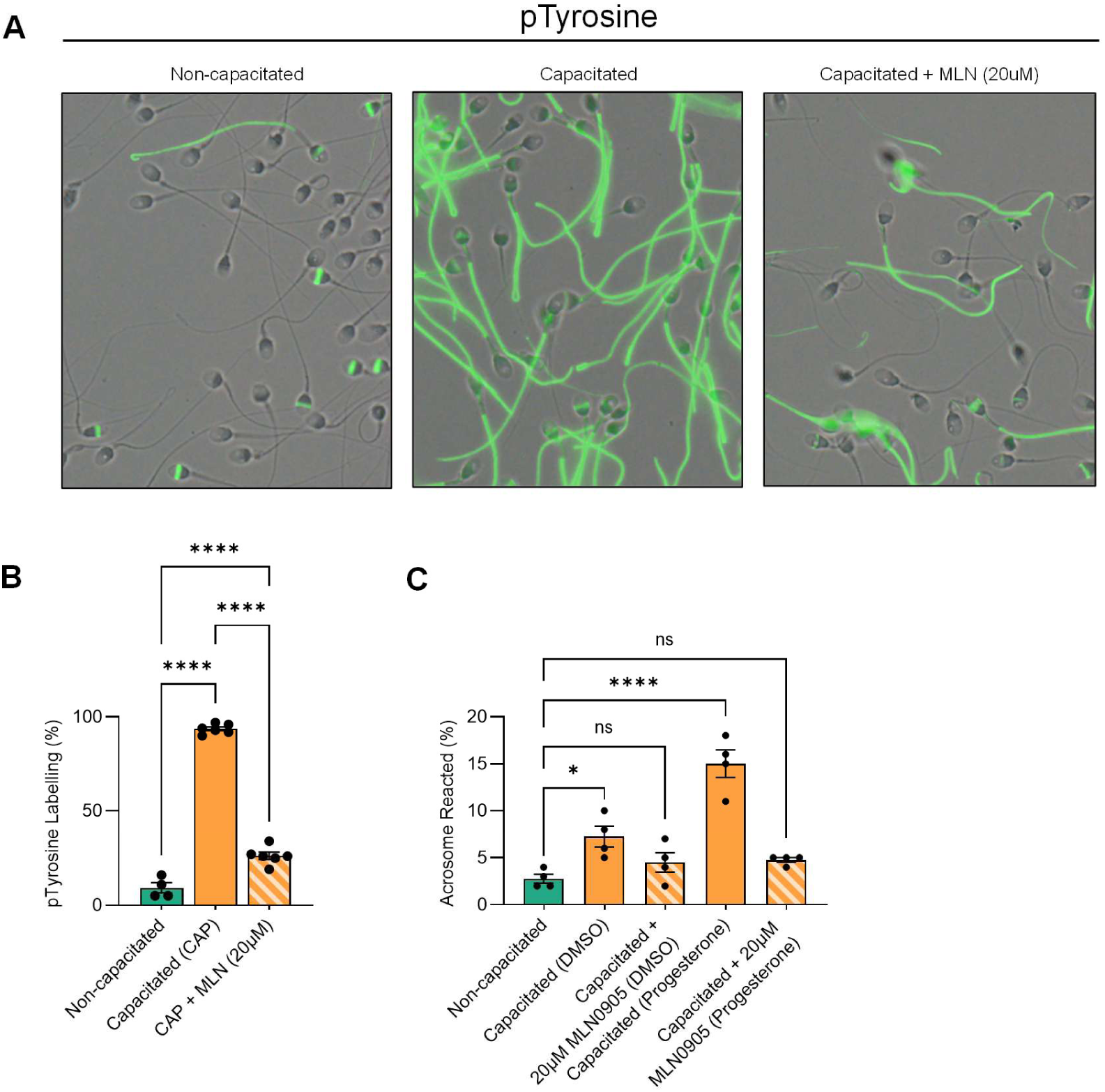
Assessment of the effect of *in vitro* PLK1 inhibition on sperm capacitation ability. Sperm were incubated in capacitating BWW with a relevant a DMSO vehicle (CAP; orange) or kinase inhibitor concentrations (orange striped) for 3 hours (37°C, 5% CO_2_), and either fixed (4% paraformaldehyde) or acrosome reacted along-side non-capacitated sperm. To assess capacitation status samples were subjected to (A) immunocytochemistry with anti-phosphotyrosine (PT66). To assess acrosomal integrity cells were labelled with FITC-PSA and acrosome reacted cells counted (B). Data were subject to one-way ANOVA with Tukey’s multiple comparisons test; * *p* ≤ 0.05, **** *p* ≤ 0.0001. All graphs contain n = 4-6 biological samples.

## DISCUSSION

Recently it has been postulated that the post meiotic windows of sperm maturation (spermiogenesis, epididymal transit and capacitation) are ideal for contraceptive targeting since they are critical steps underpinning successful fertilization that do not require manipulation of hormone secretion ^46^. In contraposition, the increasing prevalence of male infertility ^73–75^, and our inability to accurately diagnose and treat it ^76–78^, provides the impetus for investigating the shortcomings of fertilization-incompetent sperm. Therefore, the pursuit of new protein kinases that regulate capacitation, and their phosphorylated protein substrates, holds merit for the development of potential contraceptives and assisted reproductive technology (ART), as well as improving sperm selection strategies for ART; a priority area for the future of infertility research^79^. The present study was undertaken with the aim of investigating the phosphorylated proteome of ejaculated, density centrifugation enriched human sperm cells, termed throughout as non-capacitated, and capacitated human sperm. Accordingly, we have generated a comprehensive repository of phosphoproteomic data that will serve as an important resource for understanding the functional roles of human sperm proteins, their regulation by phosphorylation, and the temporal kinase-protein interactions controlling sperm activation. Our dataset will also act as a reference for future phosphoproteomic inquiry into male infertility.

Here we have curated a phosphoproteome comprising over 2,000 phosphoresidues from 843 total human sperm proteins. Inquiry into the biological processes regulated by sperm phosphoproteomic signatures indicated an enrichment for motility related conditions and functions, including asthenozoospermia, flagellar abnormalities and cell movement of sperm. This illustrates the investment required to achieve motility in these cells, adding credence to the use of phosphoproteomics to investigate the mechanistic basis of sperm motility and asthenozoospermia ^54–57^.

The second most enriched biological process from IPA was spermatogenesis, namely the terms spermatogenic impairment, spermatogenic failure, and meiosis of germ cells. Noting that our phosphoproteomic analysis was conducted on mature sperm, this likely indicates the limited annotation of sperm and fertility related proteins in broad databases like this that are usually built from somatic cell data. This notion is also emphasized by proteomic studies on mouse sperm, including those by Skerget et al. ^82^ and Skerrett-Byrne *et al.* ^6^, which demonstrate a refining of the sperm molecular landscape coinciding with the enrichment of key cellular processes including “*capacitation of sperm*” and “*penetration of zona pellucida*”, as well as “*cell movement of sperm*” featured here, during the post-testicular phases of sperm maturation. As such, we propose that the phosphoproteins curated in the mature sperm phosphoproteome described herein should not be considered exclusively in the context of previously assigned roles in spermatogenesis, but rather they should be subjected to further investigation to better appreciate the implications of their retention throughout post-testicular maturation and the functional importance linked to their phosphorylation status.

In terms of the nature of capacitation-associated phosphorylation events, we observed dynamic changes between the three phosphorylated amino acids (phosphoserine, phosphothreonine, and phosphotyrosine), including increases in phosphoresidue abundances across all three amino acids linked with *in vitro* capacitation. However, there was a noticeable disparity between non-capacitated and capacitated cells in the abundance of phosphothreonine and phosphotyrosine modified phosphopeptides, tending towards higher abundances among capacitated sperm. Notably, phosphorylated tyrosine comprised 3.0% of the capacitated sperm phosphoproteome compared to 1.8% among non-capacitated sperm, with the detection of 18 phosphotyrosine residues uniquely present following capacitation [not-detected in non-capacitated sperm]. Understandably, the very clear increase in tyrosine phosphorylation has historically excited much interest in understanding the role of tyrosine phosphorylated proteins in sperm ^37,38,64,65^. Consequently, the contribution of tyrosine phosphorylation to capacitation, and in particular motility, is well appreciated such that it is used, globally, as a marker of sperm capacitation across most mammalian species ^83–95^. Complementing this existing literature, here we have shown that there is also an enrichment for motility related proteins and functions among those targeted for threonine phosphorylation during capacitation. Ficarro *et al.* ^37^ cite the limited utility of antibodies against phosphoserine and phosphothreonine due to a lack of sensitivity towards those modifications, particularly concerning the two-dimensional gel coupled to immunoblot enrichment method they have utilized. While this may be a historically limiting factor, the sensitivity of LC-MS/MS and the improved efficiency of phosphoproteomic sample preparation has superseded the necessity of traditional gel-immunoblot enrichments in most settings and, in this way, we can now report significant upregulation in the threonine phosphorylation of 19 peptides.

Two prior phosphoproteomic studies, by Ficarro *et al.* ^37^ and Wang *et al.* ^36^, have included comparisons between non-capacitated and capacitated human sperm. Here, we extend these studies by identifying over 750 sperm (testis/male germ line) specific phosphorylation sites even though over 90% of the parent proteins had at least one previously reported phosphorylation site. Excitingly, this may be due to unique roles for phosphorylation in sperm cells, and these newly identified sperm-specific phosphorylation sites may offer novel insights into sperm biochemistry. Similarly, previous efforts to investigate post-translational modifications on a global tissue scale identified over 200 testis specific phosphorylation sites in the rat testis ^100^, and this study along with another in mice ^101^ found that testes were second only to the brain in the number of tissue specific phosphorylation sites. Further, the development of kinase-substrate interaction databases has enhanced our ability to predict kinase activity within a phosphoproteomic dataset. Here, we compared non-capacitated and capacitated human sperm phosphoproteomes and detected 20 kinases through mass spectrometry and an additional 52 kinases through *in silico* analysis with previously uncharacterized roles in sperm capacitation. Herein, we performed *in vitro* validation of a subset of the putative kinaseswe described here.

In transitioning from *in silico* analyses to *in vitro* validation, not all pharmacological inhibitors of kinases exhibited an effect on sperm parameters. This may arise if the inhibitors utilized are not suitably efficacious for sperm, if the predicted kinases are not present (a false positive prediction), or if the kinases’ role in the spermatozoon is not congruent with the parameters we investigated. However, there should also be consideration given to the importance of the sperm cell and how its essential role in species continuation may reveal why the inhibition of individual kinases may not produce a discernible impact. The sperm of eutherian mammals must undergo two sequential post-testicular maturation events, separated by an extended residency in the distal cauda epididymis during which they attain motility, and where the latter maturation event taking place in the reproductive tract of a separate member of the species ^33^. By contrast, the sperm of well-studied avian models, including the domestic fowl (Gallus domesticus) and coturnix quail (Coturnix japonica), are already motile in the testis and exhibit a comparatively truncated transit time of the epididymis which lacks epididymal specializations associated with sperm storage ^102,103^. Instead, the onus of extended storage of sperm occurs in the female reproductive tract, where sperm can reside for an extended window, in some cases lasting months ^104^. These extreme examples of post-testicular maturation occur in sperm among the amniotes, whereas marsupials and monotremes ^105–107^, and the reptilian genus including Crocodilia ^108–110^, reflect combined elements of these maturation styles. The diversity and complexity of fertilization mechanics, prominent among which is the switching on and off of cellular functions including energy metabolism and motility through phosphorylation signaling, lends itself to the establishment of redundancies as a necessity to ensure that sperm are fit to compete to fertilize the oocyte ^111^. In this way, important redundancies arise where single protein substrates may be phosphorylated by numerous kinases or where a single sperm function, such as motility or the acrosome reaction, may be governed by more than one pathway.

With this in mind, we selected a subset of kinases without previously described roles in mature sperm or capacitation for targeted pharmacological inhibition. Small molecule kinase inhibitors are heralded as a cornerstone of therapeutic discovery ^112^, and in functionally mature human sperm, where *in-vitro* culture and genetic manipulation are not possible, small molecule inhibitors are crucial to developing our understanding of sperm physiology and molecular functions. However, an important consideration when using small molecule inhibitors is their propensity for off-target effects which are generally attributed to non-specific binging or cross-talk/retroactivity ^112,113^. In this study, inhibition of AKT1 did not yield any appreciable alterations to viability, motility, or phosphotyrosine immunodetection (Figure S11A-H). In contrast, it has been observed that the phosphatidylinositol 3-kinase (PIK3) signaling pathway phosphorylates, and thus activates, AKT1 in capacitated human sperm following progesterone stimulation, and that AKT1 activation mediates sperm hyperactivation ^114^. Though our observations differ, this is likely due to the progesterone dependent nature of AKT1 activation reported by Sagare-Patil *et al.* ^114^. In non-capacitated human sperm CK2 inhibition failed to perturb viability or any of the observed motility parameters. Similarly, in capacitated human sperm CK2 inhibition failed to significantly reduce anti-phosphotyrosine labeling (immunoblotting), nor viability or motility with the exception of reducing VSL at a high concentration (50 μM CX-4945). This decrease in VSL, though significant, offers little merit for CK2 inhibition as a target for contraceptive purposes and in fact offers little evidence in favor of the function of CK2 in the mature human spermatozoon. We caution ruling out the possibility entirely, as CK2 may play a role in sperm egg interaction or other biological processes not investigated here.

Similarly, PAK1 inhibition did not perturb cell viability or motility in non-capacitating sperm cells. Under capacitating conditions, viability and motility were unaffected, however progressive motility, among other CASA quantified sperm kinematics, was significantly decreased at 50 μM NVS-PAK1-1. These data allude to a potential role for PAK1 in regulating human sperm motility during capacitation, though we note that a more efficacious inhibitor is needed to further explore this putative role. Notwithstanding this limitation, previously, PAK1 was found to be involved in actin-polymerization during capacitation in mouse sperm, demonstrated through inhibition with the specific small molecule inhibitor IPA-3 ^115^.

PAK4 has been evaluated in mouse sperm previously, wherein selective inhibition with 10 μM PF-3758309 under capacitating conditions significantly hampered actin polymerization and acrosomal exocytosis and identified that PAK4 may be responsible for phosphorylating, and consequently inhibiting, protein phosphatase Slingshot homologue 1 (SSH1) ^116^. Targeted pharmacological inhibition of PAK4 in non-capacitated human sperm in our study did not perturb viability or total motility during a 30 min incubation. Surprisingly, PAK4 inhibition resulted in a significant increase in progressive motility at both 7.5 μM and 50 μM under non-capacitating conditions in 30 min. This may indicate a role for PAK4 as a decapacitation factor or in regulating a phosphatase responsible for maintaining the inactivated state of non-capacitated human sperm. Under capacitating conditions, PAK4 inhibition did not perturb the viability or motility of human sperm, although progressive motility and other CASA kinematics were significantly decreased to levels reflective of non-capacitated sperm. Likewise, PAK4 kinase inhibition reduced tyrosine phosphorylation levels. This marked decrease in tyrosine phosphorylation, paired with the impact of PAK4 inhibition on progressive motility, indicates that PAK4 kinase is likely present and actively regulating sperm capacitation activity. One caveat is that the Group II selectivity of our inhibitor GNE 2861 means that while the inhibitor is most effective against PAK4 (5-fold and 17-fold increased selectivity compared to PAK5 and PAK6 respectively), further inhibitor and activator studies are required to eliminate the prospect of additional inhibitory effects on related protein kinases in human sperm.

Finally, PLK1 inhibition, significantly reduced progressive motility, and other sperm kinematics, under capacitating conditions. Additionally, immunoblot analysis of tyrosine phosphorylation revealed a significant reduction in tyrosine phosphorylation to levels equal to non-capacitated human sperm. Further, probing for anti-phosphotyrosine at a cellular level revealed that the prescribed anti-phosphotyrosine tail labeling associated with capacitation was inhibited in a majority of human sperm when capacitated during PLK1 inhibition. PLK1 inhibition was also sufficient to suppress both spontaneous and progesterone induced acrosome reaction rates in sperm under capacitating conditions. This line of inquiry provides strong evidence for an active role for PLK1 in regulating capacitation, the acrosome reaction, and progressive motility. PLK1 kinase has been reported to phosphorylate (and thus activate) PPP1R7, a known regulator of a sperm specific protein phosphatase (PP1γ2) ^72^. Despite this, PLK1 has not previously been the subject of investigation in human sperm. This is likely an oversight attributed to the essential nature of PLK1 in meiosis in the mouse, where conditional-germ cell deletion of PLK1 resulted in complete azoospermia with meiotic arrest, and thus the function of PLK1 has not been considered in post meiotic spermatids and mature sperm ^117^. Despite this, immunolocalization indicated the presence of PLK1 throughout the midpiece and peri-acrosomal region of mature sperm.

The PLK1 inhibitor used here, MLN0905, has been evaluated for effectiveness against a panel of 395 kinases and was identified to be selective for PLK1 ^118^. In screening, MLN0905 was more efficacious towards the inhibition of PLK1 than other kinases, though Fer tyrosine kinase (FER) was among the kinases with some potential cross-reactivity. FER has been well characterized in murine sperm ^119,120^, and a recent study in human sperm reported a role for FER as a negative regulator (suppressor) of human sperm capacitation ^118^. Specifically, following FER inhibition with E260 in human sperm under capacitating conditions, Duffey *et al.* ^118^ reported increased hyperactivated motility. This did not mirror our results using MLN0905 to inhibit PLK1 where no changes in CASA kinematics were observed that would be indicative of an altered hyperactivation pattern, such as an increase in beat cross frequency (BCF). Duffey *et al.* ^118^ further reported increased spontaneous acrosome reaction rates in human sperm under conditions of FER inhibition using E260, in contrast we reported PLK1 inhibition with MLN0905 decreased both spontaneous acrosome reactions and progesterone induced acrosome reactions. These contrasting observed results increase our conviction that MLN0905 is unlikely to be impacting FER kinase, and that PLK1 indeed is responsible for the impacts of MLN0905 in sperm function. While the definitive mode of action of PLK1 in sperm capacitation has not been elucidated here, we speculate that PLK1 is likely to be active earlier in the cascade of kinase activity and may act either in parallel to or downstream from PKA as a higher order orchestrator of sperm capacitation events. In this way PLK1, a serine/threonine kinase, could indirectly effect capacitation dependent tyrosine phosphorylation through modulation of downstream contributors to capacitation, such as tyrosine kinases, by regulating phosphatase activity (i.e. PPP1R7 and PP1γ2), and consequently perturbing progressive motility. Reconciling the role of PLK1 in sperm will be pertinent to understanding PLK1’s overall mechanism of action in addition to exploring its contributions to the regulation of sperm functional parameters not studied here, including surface remodeling events, sperm-egg interaction, and fertilization.

### Limitations of the study

Here we highlight possible caveats for the interpretation of these data and identify experiments that should be conducted to further validate our findings. The human sperm cells utilized throughout this study are from donor ejaculates, and while density gradient centrifugation is used to remove seminal plasma and contaminating cells, exposure to bicarbonate present in the seminal plasma may stimulate kinase activity, thus changing phosphorylation in contrast to quiescent epididymal sperm cells. This should be considered in the interpretation of the data beyond the scope of this manuscript where the term “non-capacitated” represents ejaculated, density centrifugation enriched human sperm cells, a convention used elsewhere for similar comparisons ^36^, and not surgically extracted epididymal sperm. It would be enlightening to perform a phosphoproteomic investigation in epididymal human sperm, establishing the temporal and spatial remodeling of both the proteome and phosphoproteome as these cells become functionally mature via epididymal maturation. Additionally, the temporal nature of phosphorylation status throughout capacitation should be examined, expanding on the two time point study convention we have adopted here. However, a low yield of cells from percutaneous epididymal sperm aspiration from the epididymis [suitable for ICSI only], and the invasive nature of the procedure, results in a paucity of such samples for human studies ^96^. Such a study may be possible in instances of epididymectomy permitting isolation of epididymal sperm or through whole tissue applications including matrix assisted laser desorption ionization mass spectrometry (MALDI-MS) ^97,98^. In seeking to understand the function and localization of the phosphoproteins identified in this study we have used GO annotations. As GO annotations are largely reliant on the localization and function of proteins in somatic cells, we advise caution in the extrapolation of this protein localization to sperm ^81^. Efforts by the Human Protein Atlas have made progress on this front, with the ‘tissue-based map of the human proteome’ project having annotated 645 human sperm proteins with validated site-specific locations ^122^. Further, in this study we induced human sperm capacitation *in vitro* using bicarbonate-based sperm capacitation media (BWW) supplemented with a phosphodiesterase inhibitor and cAMP analogue. Although this is the most widely published method of inducing human sperm capacitation for experiments *in vitro* to date ^63–65,94,95,123–128^, it does not necessarily reflect the protocols used in assisted reproduction clinics. This difference in media choice could have an impact on subtle aspects of phosphorylation signaling and this should be considered when extrapolating these data to clinical applications. Additionally, the use of the cAMP analogue may bias capacitation pathways that are dependent on protein kinase A and thus we suggest that follow up studies should track capacitation induced phosphorylation changes in bicarbonate-based capacitation media alone as well as in media used in many IVF clinics worldwide ^129^. Challenges to the paradigm of sperm capacitation have called into question tyrosine phosphorylation as a marker of fertilization competent sperm in the mouse due to (1) its transient nature when visualized in *situ* ^130^ and (2) due to observations that the sperm of tyrosine kinase FER null mice can fertilize in the absence of bulk increases in tyrosine phosphorylation ^119^. However, these experiments have not been repeated in humans. Our data adds credence to an important role for tyrosine phosphorylation in the fertilization cascade, as well as an unappreciated role for threonine phosphorylation. However, we encourage further experiments to explore the precise spatial and temporal nature of phosphorylation changes in human sperm during capacitation.

## Conclusion

Here, we provide the first proteomic evidence of site-specific phosphorylation in 755 phosphoproteins and report a curated library of putative capacitation kinases that can now be investigated by the field as targets for the regulation of fertility. Contributing to such investigations, we have provided evidence that PAK4 has a putative functional role as a decapacitation factor in human sperm limiting the induction of capacitation, and that PAK1, PAK4 and PLK1 may contribute to the establishment of progressive motility in capacitating human sperm. Moreover, we have demonstrated that PLK1 contributes to the establishment of widespread protein tyrosine phosphorylation and critical pre-fertilization events (e.g the acrosome reaction) in human sperm positively associated with fertilization capacity. To further establish their contributions to male fertility, each of these kinases requires the application of technology to identify their interacting partners, activation signals, phosphorylated substrates, and to determine redundancies in their function. This study delivers the largest depth in the human sperm phosphoproteome and predicted sperm kinome to date, greatly expanding our capacity to develop sperm-focused contraceptives and therapies for sperm dysfunction.

### TABLE 3: **KEY RESOURCES TABLE**

| REAGENT OR RESOURCE | SOURCE | IDENTIFIER |
| --- | --- | --- |
| Antibodies |  |  |
| Phosphoserine (pSerine) | Sigma Aldrich | Cat# P5747 ; RRID:AB_477376 |
| Phosphothreonine (pThreonine) | Sigma Aldrich | Cat# P6623; RRID:AB_477393 |
| Phosphotyrosine-HRP | Sigma Aldrich | Cat# A5964; RRID:AB_476748 |
| Phosphotyrosine (pTyrosine) | Sigma Aldrich | Cat# P5872; RRID:AB_1079609 |
| GAPDH | Sigma Aldrich | Cat# G9545; RRID:AB_796208 |
| α-Tubulin | Cell Signaling Technology | Cat# 2125; RRID:AB_2619646 |
| Goat-α-mouse HRP | Dako Denmark | Cat# P0447; RRID:AB_2617137 |
| Goat-α-rabbit-HRP | Dako Denmark | Cat# P0448; RRID:AB_2617138 |
| Goat-α-mouse 488 | Invitrogen | Cat# A-21202; RRID:AB_141607 |
| PLK1 | Sigma Aldrich | Cat# WH0005347M1-100UG; RRID:AB_1843038 |
| α-Tubulin | Abcam | Cat# AB4074; RRID:AB_2288001 |
| Glycerokinase | Proteintech | Cat# 13360-1-AP; RRID:AB_1167391 |
| AKAP3 | Proteintech | Cat# 13907-1-AP; RRID:AB_2273887 |
| Donkey-α-rabbit 555 | Invitrogen | Cat# A31572; RRID:AB_162543 |
| Bacterial and virus strains |  |  |
| Biological samples |  |  |
| Chemicals, peptides, and recombinant proteins |  |  |
| Critical commercial assays |  |  |
| Pierce™ BCA Protein Assay Kit | Thermo Fisher Scientific | Cat#23225 |
| Trypsin/Lys-C Mix, Mass Spec Grade | Promega Corp. | V5072 |
| DC Protein Assay (detergent compatible) | Bio-Rad | 5000111 |
| Deposited data |  |  |
| Reference Proteome – *Homo sapiens* | UniProt | UP000005640; RRID:SCR_004426 |
| Human sperm – phosphoproteome data | This paper | PXD066153; RRID:SCR_003411 |
| Experimental models: Cell lines |  |  |
| Experimental models: Organisms/strains |  |  |
| Oligonucleotides |  |  |
| Recombinant DNA |  |  |
| Software and algorithms |  |  |
| Proteome Discoverer 2.5 | Thermo Fisher Scientific | RRID:SCR_014477 |
| Perseus (version 1.6.14.0) | (Tyanova et al., 2016) | RRID:SCR_015753 |
| Phosphomatics | (Leeming et. Al.,2021) | https://phosphomatics.com/ |
| RoKAI | (Yilmaz et. Al., 2021) | https://rokai.io/ |
| Ingenuity Pathway Analysis | Qiagen | RRID:SCR_008653 |
| KinMap | Cell Signaling Technology, Inc. | http://www.kinhub.org/kinmap/ |
| GraphPad Prism (version 9.5.1) | Graph pad | RRID:SCR_002798 |
| ImageJ 1.53 | NIH – Public Domain | RRID:SCR_003070 |
| Other |  |  |
| Leja 20µm fixed depth, four chamber slides | IVM technologies | Ref. 025107-025108 |
| CX-4945 | Jomar Life Research (Melbourne, Australia) | No.S2248; CAS No. 1009820-21-6 |
| NVS-Pak1-1 | Jomar Life Research (Melbourne, Australia) | No.S8264; CAS No. 1783816-74-9 |
| GNE 2861 | Jomar Life Research (Melbourne, Australia) | No.E2380; CAS No. 1394121-05-1 |
| MLN0905 | Jomar Life Research (Melbourne, Australia) | No.S2898; CAS No. 1228960-69-7 |
| MK-2206-2HCL | Jomar Life Research (Melbourne, Australia) | No.S1078; CAS No. 1032350-13-2 |

### RESOURCE AVAILABILITY

#### Lead contact

Further information and requests for resources and reagents should be directed to and will be fulfilled by the lead contact, Elizabeth G Bromfield.

## Materials availability

All reagents were purchased from Merck and were of molecular biology or research quality unless otherwise specified. Kinase inhibitors were purchased from Jomar Life Research (Melbourne, Australia). See Table 3 for full resource details including reference data and all software versions used.

## Data and code availability

- The mass spectrometry proteomics data have been deposited to the ProteomeXchange Consortium (http://proteomecentral.proteomexchange.org) via the PRIDE partner repository (Perez-Riverol et al. ^131^) with the dataset identifier PXD066153.

`o` PRIDE Reviewer account details:
`o` Username:
`o` Password: ItVuW3cVPTmG
- Original western blot images will be deposited at Mendeley and will be publicly available as of the date of publication. Microscopy data reported in this paper will be shared by the lead contact upon request.
- Code: this paper did not report original code.
- Any additional information required to reanalyze the data reported in this paper is available from the lead contact upon request.

### EXPERIMENTAL MODEL AND STUDY PARTICIPANT DETAILS

#### Ethical approval

The experiments described in this study were conducted using human semen samples obtained with informed consent from a panel of healthy donors. Semen parameters were assessed for each sample obtained and were conducted in accordance with criteria set in the sixth WHO laboratory manual for the examination and processing of human semen ^132^. All studies were performed in accordance with the University of Newcastle’s Human Ethics Committee guidelines (Approval No. H-2013-0319). A total of 46 samples were included in this study, from 17 donors, as detailed in Table S4.

### METHOD DETAILS

#### Preparation of human sperm

The isolation of unequivocably non-capacitated cells can only be achieved through surgical collection of cauda-epididymal sperm cells; however, a low yield of cells from percutaneous epididymal sperm aspiration from the epididymis [suitable for ICSI only], and the invasive nature of the procedure, results in a paucity of such samples for human studies ^96^. Such a study may be possible in instances of epididymectomy permitting isolation of epididymal sperm or through whole tissue applications including matrix assisted laser desorption ionization mass spectrometry (MALDI-MS) ^97,98^. In lieu of surgically collected sperm, for the purpose of this study, ejaculated human sperm prepared via density gradient centrifugation in bicarbonate-depleted media are termed non-capacitated. Here, non-capacitated sperm represent the most immature state authors could obtain sperm cells in, which are contrasted to sperm cells driven to capacitate in bicarbonate enriched media supplemented as described herein. Ejaculated human sperm samples were prepared following a 30 min liquefaction period, with semen analysis performed in accordance with WHO criteria and presented in Table S4. Semen samples were layered over a discontinuous Percoll density gradient (comprising 40% and 80% Percoll suspensions) and fractionated by centrifugation at 500 × g for 30 min ^29,133^. To isolate the enriched sperm pellet at the base of the 80% Percoll fraction, nominally referred to as ‘good quality’ sperm ^134^, the overlying Percoll was removed. Sperm were resuspended in non-capacitating (NaHCO_3_ free) Biggers, Whitten and Whittingham (BWW) medium ^135^ and washed by centrifugation at 500 × *g* for 15 min. BWW consisted of 120 mM NaCl, 4.6 mM KCl, 1.7 mM CaCl_2_2H_2_O, 1.2 mM KH_2_PO_4_, 1.2 mM MgSO_4_7H_2_O, 5.6 mM D-glucose, 0.27 mM sodium pyruvate, 44 mM sodium lactate, 5 U/ml penicillin, 5 mg/ml streptomycin, 20 mM HEPES buffer, 1 mg/ml PVA (substituted for BSA; osmolarity of 290-310 mOsm/kg) ^29^. A subset of cells was assessed for motility, viability using eosin [0.5% (w/v) eosin B, 0.9% (w/v) NaCl, in H_2_O], and concentration. Sperm were then collected as non-capacitated samples or, following a centrifugation event (500 × g, 3 min), resuspended in capacitating BWW (10 × 10^6^ cells/mL) and capacitated for 3 hours at 37°C under a 5% CO_2_/95% air atmosphere with a gentle inversion every 30 min. Capacitating BWW was enriched with 25 mM NaHCO_3_ and supplemented with 3 mM pentoxifylline (ptx) and 5 mM dibutyryl cyclic adenosine monophosphate (dbcAMP), a membrane permeable cyclic AMP analogue ^63,64,136^. Following capacitation, a subset of cells was assessed for motility and viability.

#### Phosphoproteomic sample preparation (EasyPhos)

Samples were processed using an established, high throughput method termed EasyPhos ^6,60,68,137–139^. Briefly, each sample (∼50 × 10^6^ sperm) was resuspended in ice cold lysis buffer [4% (w/v) sodium deoxycholate (SDC), 100 mM Tris-HCl (pH 8.5)], and immediately heated (95°C, 8 min) to inactivate endogenous proteases and phosphatases. Samples were sonicated on ice (4 × 30 s cycle; 10 s 100% output power, 20 s rest on ice), and an aliquot of homogenate was taken to determine protein concentration using a bicinchoninic acid assay (BCA; Pierce BCA Protein Assay Kit, Thermo Fisher Scientific, Waltham, MA). Equal starting amounts of protein (120 μg) were diluted to 270 μL in lysis buffer in a 2 mL deep well plate. Samples were simultaneously reduced and alkylated [10 mM Tris(2-carboxyethyl)phosphine hydrochloride, 40 mM 2-chloroacetemide], using a Thermomixer (Eppendorf; Hamburg, Germany) in which samples were incubated for 5 min at 45°C (1,500 rpm). Enzymatic digestion was achieved using trypsin/Lys-C mixture (V5072, Promega) in a 1:30 (w/w) enzyme-to-substrate ratio and incubated in a Thermomixer overnight (37°C, 1,500 rpm). Samples were sequentially diluted and shaken in isopropanol [3:4 (v/v) ratio, 30 s, 1,500 rpm], and phosphopeptide enrichment buffer [30 s, 1,500 rpm; 48% (v/v) trifluoroacetic acid (TFA), 8 mM KH_2_PO_4_]. Phosphopeptide enrichment was achieved using 5 mg titanium dioxide (TiO_2_) beads (Titansphere Phos-TiO bulk, 20 μm, GL Sciences, Tokyo, Japan) per peptide digest, incubated shaking (2,000 rpm) for 5 min at 40°C. Titanium dioxide beads were pelleted (2,000 × g, 1 min), proteome-supernatant removed (and stored), and TiO_2_-bound phosphopeptides washed 5 times in phosphopeptide wash buffer [5% (v/v) trifluoracetic acid (TFA), 60% (v/v) isopropanol], pelleting and discarding spent solution between washes. Washed, TiO_2_-bound phosphopeptides were resuspended in transfer buffer [0.1% (v/v) TFA, 60% (v/v) isopropanol], transferred on top of C8 (3 M Empore) two-layer StageTips ^140^, and centrifuged until dry (1,500 × g, 8 min) using a custom 3D-printed StageTip adapter ^6,141^.

Phosphopeptides were eluted in elution buffer [5% (v/v) NH_4_OH, 32% (v/v) acetonitrile] by centrifugation (1,500 × g, 3 min), and repeated to ensure complete elution of phosphopeptides. For desalting, eluates were vacuum concentrated (∼20 μL volume), diluted 1:5 (v/v) in 1% (v/v) TFA in isopropanol, transferred to styrenedivinylbenzene-reverse phase sulfonated (SDB-RPS) StageTips ^140^, and centrifuged (1,500 × g, 8 min). Sequentially, phosphopeptide-StageTips were washed in 1 % (v/v) TFA isopropanol (centrifuged 1,500 × g, 8 min) and 0.2% (v/v) TFA, 5% (v/v) acetonitrile (centrifuged 1,500 × g, 5 min). Desalted phosphopeptides were eluted [20 μL of 25% (v/v) NH_4_OH, 4 mL 60% (v/v) acetonitrile] by centrifugation (1,500 × g, 3 min) and dried by vacuum concentration. Phosphopeptides were re-suspended in MS loading buffer [2% (v/v) acetonitrile, 0.3% (v/v) TFA], and purified sperm phosphopeptides were analyzed by high resolution nano liquid chromatography tandem mass spectrometry (nLC-MS/MS).

#### nLC-MS/MS analysis

Reverse phase nLC-MS/MS was performed using an Orbitrap Exploris 480 MS coupled to a Dionex UltiMate 3000RSLC nano high-performance liquid chromatography system (Thermo Fisher Scientific). Peptide separation was then achieved using an EASY-Spray PepMap C18 75 µm x 25 cm, employing a linear acetonitrile gradient over 120 min (2–35%, 96 min; 35–95%, 4 min; 95–2%, 20 min). Full MS/data dependent acquisition MS/MS mode was utilized on Xcalibur (Thermo Fisher Scientific; version 4.2.47) with the Orbitrap mass analyzer set at a resolution of 60,000, to acquire full MS with a m/z range of 350-1500, incorporating a normalized automatic gain control target of 300% and maximum fill times of 100 ms. The 20 most multiply charged precursors were selected for higher-energy collision dissociation fragmentation with a collisional energy of 30 and 36 %. MS/MS fragments were measured at an Orbitrap resolution of 15,000 using standard mode for automatic gain control target and maximum fill time set to automatic.

#### Kinase inhibition: cell viability and computer-assisted sperm analysis (CASA)

A dose response was performed to examine initial cytotoxicity for several kinase inhibitors used in this study. The specific kinases that were inhibited were, Casein kinase II subunit alpha (CK2) with CX-4945 (Silmitasertib; 1 nM to 50 μM), p21-activated kinase 1 (PAK1) with NVS-PAK1-1 (5 nM to 50 μM), p21-activated kinase 4 (PAK4) with GNE 2861 (7.5 nM to 50 μM), and Polo-like kinase 1 (PLK1) with MLN0905 (2 nM to 20 μM). Inhibitor concentrations were chosen to reflect log scales of the IC50 values reported in cell free kinase assays performed by the manufacturers. Accompanying DMSO vehicle controls were used, at concentrations <1:1000, to mirror the highest inhibitor concentrations. Where samples were prepared for functional studies, sub-maximal inhibitor concentrations were selected, when required, to maintain DMSO-vehicle concentrations at known sub-lethal levels for sperm cells. Human sperm were diluted to 10 × 10^6^ sperm/mL in non-capacitating BWW spiked with the respective kinase inhibitors for a pre-capacitation incubation (30 min, 37°C). A subset of cells were assessed for viability (using eosin as above) and motility using computer-assisted sperm analysis (CASA) or, following centrifugation (500 × g, 3 min), were resuspended in capacitating BWW (10 × 10^6^ cells/mL) spiked with the respective kinase inhibitors and capacitated for 3 hours [in the presence of the inhibitors] at 37°C under a 5% CO_2_/95% air atmosphere with a gentle inversion every 30 min. Following capacitation, a subset of cells was assessed for viability (eosin) and motility (CASA). Kinase inhibitor treated sperm were then snap frozen in liquid nitrogen for SDS-PAGE and immunoblotting experiments, or fixed using 4% paraformaldehyde for immunolabeling, or assessed for acrosomal exocytosis.

CASA was performed using an IVOS II (Hamilton Thorne, Danvers, MA, USA) to assess sperm motility parameters. For motility assessment a total of 3 μL of sperm from each sample was loaded into one chamber of a 20 μm deep four-chambered slide (Leja; Gytech PTY Ltd, Australia). A minimum of 200 cells and five fields were analyzed per sample using a negative phase-contrast optic, a recording rate of 60 frames/s, and a stage temperature of 37°C. The following parameters were used: progressive average path velocity (VAP) threshold of 50 μm/s, slow (static) cell VAP threshold of 20 μm/s, slow (static) cell velocity (VSL) threshold of 0 μm/s, and threshold straightness (STR) of 75%. Cells exhibiting a VAP of ≥ 50 μm/s and an STR of ≥ 80% were considered progressive.

*Sodium dodecyl sulphate-polyacrylamide gel electrophoresis (SDS-PAGE) and immunoblotting* Protein was extracted from cells via boiling in sodium dodecyl sulphate (SDS) extraction buffer [0.375 M Tris pH 6.8, 2% (w/v) SDS, 10% (w/v) sucrose, protease inhibitor cocktail] at 100°C for 5 min. A cleared protein lysate was obtained following centrifugation (17 000 × g, 10 min, 4°C) to remove insoluble material, and supernatant proteins were quantified using a DC protein assay (Bio-Rad Laboratories, Hercules, CA). SDS-PAGE was performed as described previously^142^. Briefly, equivalent amounts of protein were boiled in SDS-PAGE sample buffer [2% (v/v) β-mercaptoethanol, 2% (w/v) SDS, and 10% (w/v) sucrose in 0.375 M Tris, pH 6.8, with bromophenol blue] at 100°C for 5 min. Samples were resolved by SDS-PAGE (150 V, 1 hour) before transfer to nitrocellulose membranes (350 mA, 1 hour). Membranes were subsequently blocked in appropriate blocking solution [3% (w/v) BSA/Tris-buffered saline containing 0.1% (v/v) polyoxyethylenesorbitan monolaurate (Tween 20; TBS-T). Immunolabeling of protein targets was accomplished by incubation with primary antibodies (Table S5), 3 × 10 min wash in TBS-T, and [where not directly conjugated to horseradish peroxidase (HRP)] sequential incubation was performed with appropriate HRP-conjugated secondary antibodies diluted in 1% (w/v) BSA/TBS-T. For antibody specific conditions see Table S5. Following secondary antibody incubation, membranes were again washed (3 × 10 min in TBS-T) and developed using an enhanced chemiluminescence detection kit as per the manufacturer’s recommendations (GE Healthcare, Silverwater, Australia) and viewed using a LAS4000 imaging system (Raytek Scientific Ltd., Sheffield, United Kingdom). Stripping of immunoblots and re-probing with anti-GAPDH (G9545) or anti-α-Tubulin (2125) was performed to calculate protein band intensity relative to the loading control (anti-GAPDH/anti-α-Tubulin), determined by densitometry using ImageJ software (ImageJ version 1.53) ^143^.

#### Immunocytochemistry

Non-capacitated and capacitated human sperm were fixed in 4% paraformaldehyde and washed three times with 0.05 M glycine in phosphate-buffered saline (PBS). Cells were pipetted onto poly-L-lysine-coated glass coverslips, allowed to settle overnight (4°C), and permeabilized using 0.2% Triton X-100 for phospho-antibodies, and 100% methanol 10 min for PLK1 and PLK1 co-labelled samples, as previously described ^29,136^. Non-specific binding was blocked with 3% (w/v) BSA/PBS for 1 hour at 37°C in a humid chamber. In the case of anti-phosphotyrosine (P5872), blocking consisted of 3% (w/v) BSA/PBS supplemented with an additional 10% (w/v) Normal Horse Serum (NHS; H1270) for 1 hour at 37°C in a humid chamber. All coverslips were then sequentially incubated in relevant primary antibodies (Table S5) overnight at 4°C, washed 3 × 5 min in PBS, and incubated with appropriate fluorescently conjugated secondary antibodies in 1% (w/v) BSA/PBS for 1 hour at 37°C to immunolabel target proteins. Where required, samples were counterstained with DAPI, 2 min ambient temp, and washed 3 × 5 min in PBS. Coverslips were washed in PBS (3 × 5 mins) before mounting in 10% Mowiol 4-88 (Merck Milipore, Darmstadt, Germany) with 30% glycerol in 0.2 M Tris (pH 8.5) and 2.5% 1,4-diazabicyclo-(2.2.2)-octane. Cell labeling patterns were examined with a Zeiss Axio A.2 fluorescence microscope (Carl Zeiss Micro Imaging GmbH, Jena, Thuringia, Germany). Immunolabeling counts were performed at 100 × magnification on 100 cells per biological replicate and treatment.

#### Assessment of acrosomal exocytosis

Following capacitation, sperm (10 × 10^6^ cells/mL) were incubated in capacitating BWW containing a final concentration of either 15 μM progesterone, 2.5 μM calcium ionophore A23187, dimethyl sulfoxide (DMSO; 1:4000 v/v) as a vehicle control ^63,65^ Cells were incubated for either 15 min (DMSO and progesterone) or 30 min (A23187) at 37°C to induce the acrosome reaction. To simultaneously assess cell viability, samples (including a non-capacitated control at time-zero) were transferred to pre-warmed hypo-osmotic swelling medium (HOS; 25 mM sodium citrate, 75 mM fructose) and incubated at 37°C for a further 1 hour, gently inverting each 15 min, as described previously ^38,144,145^. Post HOS incubation, cells were fixed in 4% paraformaldehyde, washed three times and then resuspended in 0.05 M glycine in PBS. A 10 μL aliquot of the fixed sperm suspension was then air dried onto Trajan 12-well diagnostic slides. Immunofluorescent labeling of glycoconjugates residing in the sperm-outer acrosomal membrane was then utilized for assessment of acrosomal exocytosis using fluorescein isothiocyanate-conjugated P*isum sativum* agglutinin (FITC-PSA), as previously detailed ^146^. Briefly, sperm were permeabilized with dropwise application of ice-cold methanol to wells and incubated for 10 min at ambient temperature in a humid chamber. Slides were then incubated with 20 μg/mL FITC-PSA (concentration determined via serial dilution, L0770-2mg, Sigma Aldrich) in PBS for 30 min, 37°C, under humid conditions, washed (3 × PBS), and mounted for imaging with 10% Mowiol 4-88 (Merck Milipoire, Darmstadt, Germany) with 30% glycerol in 0,2 M Tris (pH 8.5) and 2.5% 1,4-diazabicyclo-(2.2.2)-octane (DABCO). Samples were examined with a Zeiss Axio A.2 fluorescence microscope (Carl Zeiss Micro Imaging GmbH, Jena, Thuringia, Germany) for detection of fluorescent-labeling patterns (Ex. 474nm, Em. ∼524nm) in viable sperm.

### QUANTIFICATION AND STATISTICAL ANALYSIS

#### Proteomic data processing and in-silico analysis

Database searching of raw files was performed using Proteome Discoverer 2.5 (Thermo Fisher Scientific), in line with previous studies ^6,68,147–149^. SEQUEST HT was used to search against the UniProt *Homo sapiens* database (42,280 sequences, downloaded 14^th^ February 2022). Database searching parameters included up to two missed cleavages, a precursor mass tolerance set to 10 ppm and fragment mass tolerance of 0.02 Da. Trypsin was designated as the digestion enzyme and cysteine carbamidomethylation was set as a fixed modification while phosphorylation (serine (S), threonine (T), tyrosine (Y)) was designated as a dynamic modification. Interrogation of the corresponding reversed database was also performed to evaluate the false discovery rate (FDR) of peptide identification using Percolator ^150^ based on q-values, which were estimated from the target-decoy search approach. To filter out target peptide spectrum matches over the decoy-peptide spectrum matches, a fixed FDR of 1% was set at the peptide level. Normalization and fold change calculation were based on label free quantitation (Minora Feature Detector, Feature Mapper, and Precursor Ions Quantifier ^149^ in Proteome Discoverer 2.5. The peptide list was exported from Proteome Discoverer 2.5 as a Microsoft Excel file and further refined to include peptides with a quantitative value in a minimum four of five biological replicates in at least one treatment condition. The refined peptide list was loaded into Perseus (version 1.6.14.0)^151^ to calculate statistical significance (*t*-test) comparative testing, and for the generation of scatter plots, principal component analysis, and heatmaps. Basic data handling, if not otherwise specified, was conducted using Microsoft Excel 365 (version 2211, Microsoft Corporation, Redmond, WA) and GraphPad Prism version 9.5.1 for Windows (GraphPad software; San Diego, CA).

For phosphopeptides with quantitative values in at least four of five replicates, the corresponding UniProt accession and transformed ratios were analyzed using Ingenuity Pathways Analysis software (IPA®, Qiagen, Hilden, Germany) as previously described ^6,68,148,149^. Analysis on the basis of predicted subcellular location and classification (other excluded), ‘*molecular and cellular functions*’ and ‘*physiological system development and function*’ were carried out using the IPA *p*-value enrichment score ^62^ for non-capacitated and capacitated sperm phosphoproteomes. We applied a stringency criterion of -log_10_ *p*-value ≥ 1.3 (i.e., *p*-value ≤ 0.05) for each group to elucidate the most significant changes in our analysis. Analysis of biological processes including canonical pathways, upstream regulators, and disease and function analyses were assessed using *p*-value, an enrichment measurement of the overlapping proteins from the dataset in a particular pathway, function or disease ^62^.

To provide further biological context to the phosphoproteomes, we loaded the normalized abundances of phosphorylated residues (1,086 serine, 97 threonine, and 47 tyrosine residues) into Phosphomatics ^61^ (uploaded 5^th^ April 2023), an interactive substrate-kinase mapping tool for global phosphoproteomic data. Where no value was present, imputation of the lowest value of the site from the relevant treatment group was performed. Where more than one abundance value was available for a given amino acid [due to multiply phosphorylated peptides or overlapping peptide sequences], only the single greatest fold change was retained regardless of directionality. Abundances were Log_2_ transformed and database searches were run against PhosphoSitePlus ^67^. Here, we represent kinase-substrate interactions as the site abundances for capacitated sperm relative to the mean abundance for non-capacitated sperm, Log_2_ transformed. Similarly, Robust Inference of Kinase Activity (RoKAI) ^152^ was employed to derive relationships between quantitative phosphoresidue data and upstream kinases (uploaded 3^rd^ April 2023). RoKAI analysis was performed “raw” without further normalization of data, using only residues in the dataset upload and no “sites in the functional neighborhood”, and through interrogating the PhosphoSitePlus “kinase substrate dataset”.

#### Immunoblot densitometry

ImageJ software (ImageJ version 1.53) ^143^ was used to perform densitometric analysis of all immunoblots. Briefly, gel images were converted to grey scale (32-bit) before each lane was labelled and the “Analyze > Gels > Plot Lanes” tool was used to visualize band intensity. Either the entire lane or the correct molecular weight bands of interest were selected. The “Gel Analysis > Plot lanes”. The “Wand” tool was subsequently used to generate the densitometric volume for each band/lane of interest. This process was repeated with the corresponding GAPDH or α-Tubulin immunoblot image to enable the volume of the band of interest to be normalized against that of the comparable loading control band volume for each lane. Relative densitometric volumes are presented as the mean of at least 3 biological replicates ± SEM.

#### Statistics

Proteomic analyses were performed using pooled (×2 donors) human sperm cell populations of 100 × 10^6^ cells. Proteomics samples were divided evenly in two and one portion was induced to undergo capacitation *in* vitro as described above (n = 5 biological replicates). All subsequent experiments were replicated at least 3 times with independent biological replicates comprising pooled sperm cells from at least two donors as a concentration of 10 × 10^6^ cells/ml media. After testing data for normality (Shapiro-Wilk normality test), statistical analyses were performed using either a two-tailed, unpaired Student *t*-test using Microsoft Excel (Microsoft Office 365), a comparative analysis *t*-test using Perseus (version 1.6.14.0), or a one-way ANOVA, with multiple comparisons testing as appropriate, using GraphPad Prism (version 9.5.1). In all cases, differences were considered significant for *p* ≤ 0.05, and significance level was denoted such that *p* ≤ 0.05 (*), *p* ≤ 0.01 (**), *p* ≤ 0.001 (***), and *p* ≤ 0.0001 (****) and details on the specific statistical analyses used feature in relevant figure legends. The number of biological replicates used in each experiment are presented in figure captions. Graphical data were prepared using GraphPad Prism (version 9.5.1) and are presented as mean values ±SEM.

### ADDITIONAL RESOURCES

N/A

## ACKNOWLEDGMENTS

The authors gratefully acknowledge helpful discussions regarding the data in this manuscript from Prof. Moira O’Bryan, Joseph Nguyen, Dr. Brendan Houston, Dr. Michael Leeming, Assoc. Prof. Tzviya Zeev Ben Mordehai, Dr. Miguel Leung and Shannon Smyth. We would also like to acknowledge the Charles Perkins Centre (Sydney Mass Spectrometry), The University of Sydney, Nathan Smith at The University of Newcastle and Dr. Sean Humphrey for providing 3D printed parts for EasyPhos and advice on protocol development. This research was supported by an Australian Research Council Discovery Early Career Researcher Award (DECRA; DE210100103) awarded to E.G.B, a National Health and Medical Research Council of Australia (NHMRC) Ideas grant to B.N: APP1147932 and NHMRC Senior Research Fellowship (APP1154837) and a National Health and Medical Research Council of Australia (NHMRC) Emerging Leadership Fellowship (APP2034392) awarded to D.A.S.B.

## AUTHOR CONTRIBUTIONS

Conceptualization, E.G.B, D.A.S.B, N.D.B, B.N.; methodology, N.D.B, D.A.S.B., A.L.A., E.G.B.,

H.C.M, K.M., H.M.H, J.E.S.; Investigation, N.B., D.A.S.B, A.L.A, G.E.B,; writing – original draft, N.B.; writing – review & editing, E.G.B, S.D.R, J.E.S, B.N, D.A.S.B, R.J.A, G.E.B, H.C.M.; funding acquisition, E.G.B. and B.N.; resources, A.L.A, H.M.H, H.C.M, K.M, D.A.S.B.; supervision, E.G.B, S.D.R, B.N, J.E.S.

## DECLARATION OF GENERATIVE AI AND AI-ASSISTED TECHNOLOGIES

Generative AI and AI-assisted technologies were not used in the production of this manuscript.

---

**Supplementary Figure 1:**
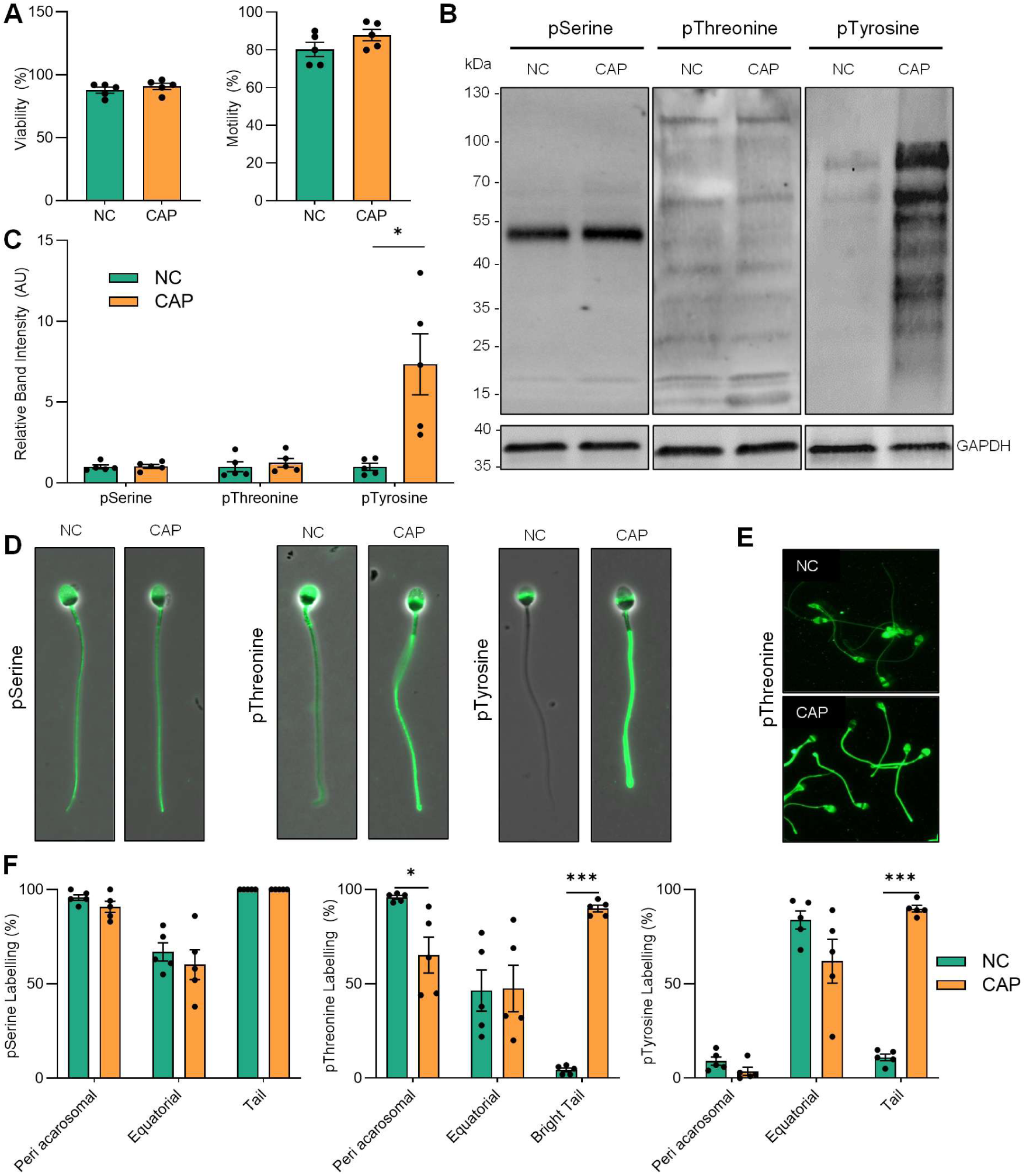
Validation of capacitation in human sperm samples used for EasyPhos and mass spectrometric analysis. (A) Viability (eosin) and motility. (B) Western blots, (C) densitometry, and (D and E) immunocytochemistry for anti-phospho antibodies. (F) Phospho-antibody labeling pattern counts. A-F n = 5 biological samples for all graphs. Data were subject to t-test; * *p* ≤ 0.05, *** *p* ≤ 0.001.

**Supplementary Figure 2:**
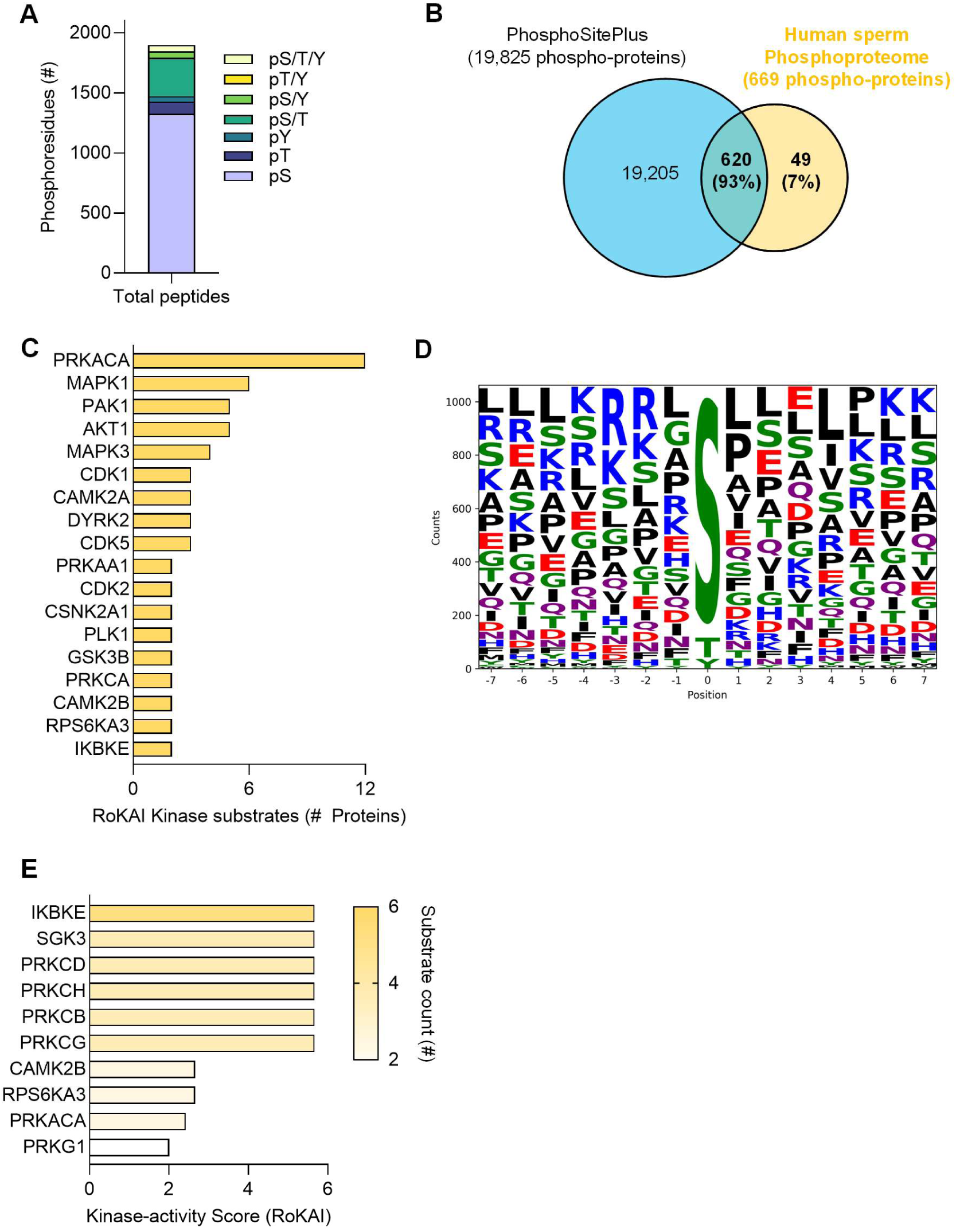
Further *in-silico* analysis of the human sperm phosphoproteome. (A) Bar graph depicting the distribution of Serine (pS), Threonine (pT), and Tyrosine (pY) phosphorylation between non-capacitated (NC) and capacitated (CAP) populations; including ambiguous sites. (B) Venn diagram comparing common and unique phospho-proteins identified in this study (yellow) and previously reported (PhosphoSitePlus; blue). (C) Bar graph of kinases with ≥2 protein substrates detected in at least one population (non-capacitated sperm and/or capacitated sperm; RoKAI accessed 15/03/2023). (D) Analysis of residues ±7 amino acids from detected phosphorylation sites. (E) Bar graph of kinases predicted (*in silico analysis*) to be active in sperm based on significantly upregulated phosphorylation (RoKAI accessed 15/03/2023).

**Supplementary Figure 3:**
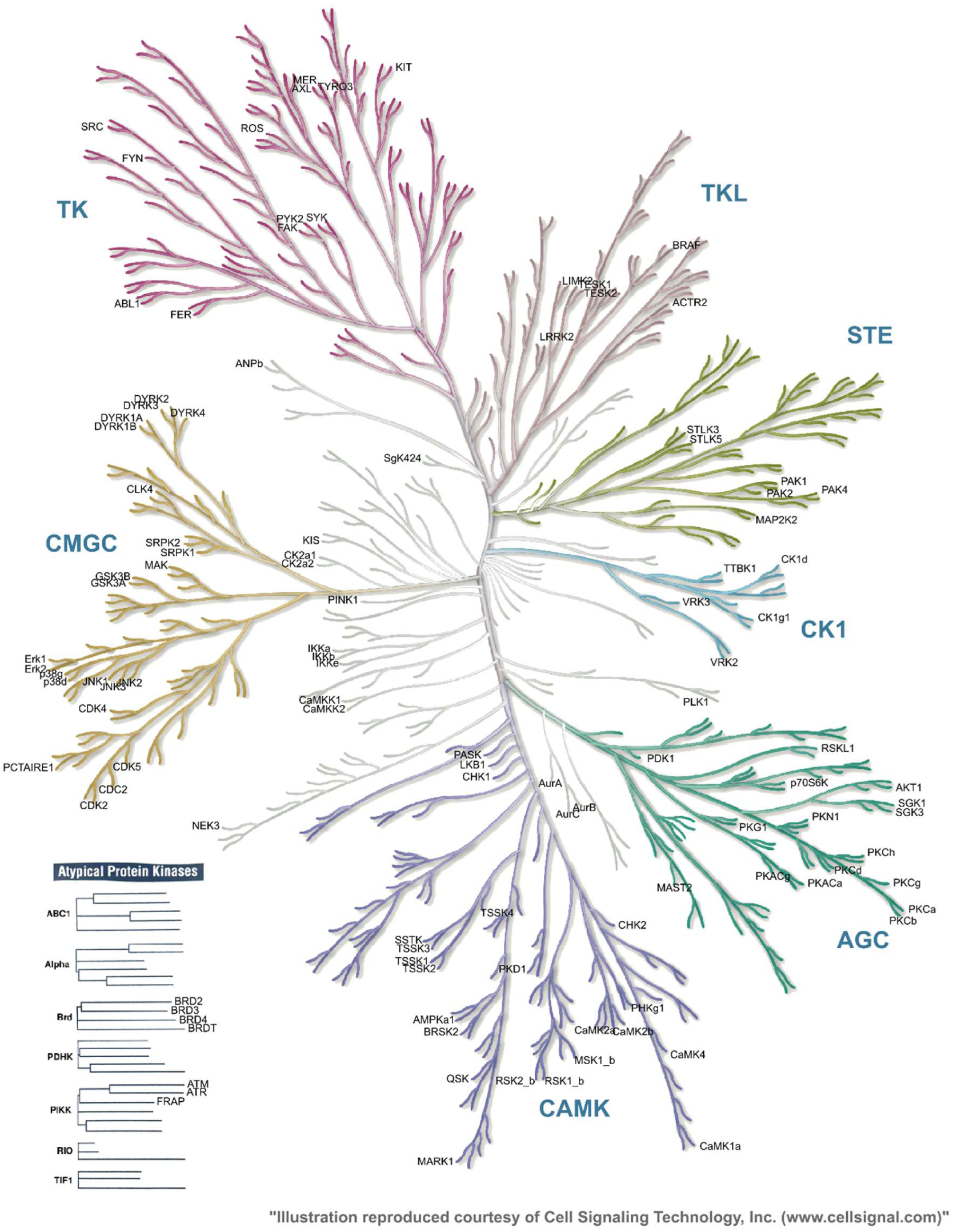
The human sperm kinome, annotated. (Cell Signaling Technology, Inc.)

**Supplementary Figure 4:**
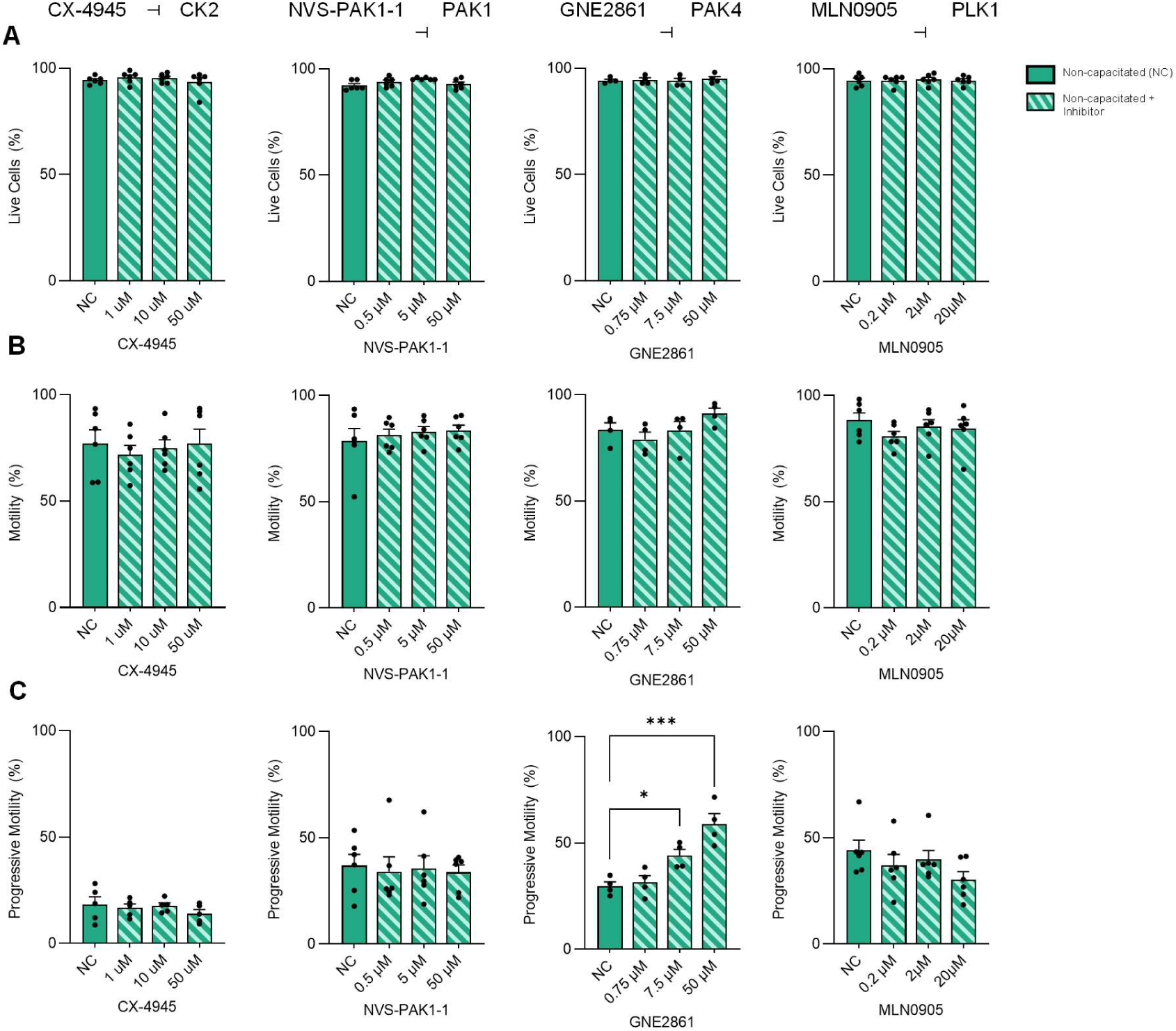
Assessment of the effect of *in vitro* kinase inhibition on sperm viability and motility parameters in non-capacitating conditions. Sperm suspensions in non-capacitating BWW were “spiked” with kinase inhibitors to relevant concentrations and pre-incubated for 30 minutes (NC; green) and subjected to (A) viability (eosin) and (B-C) motility assessment (CASA). Data were subject to a non-parametric one-way ANOVA with Dunnett’s multiple comparisons test; * *p* ≤ 0.05, *** *p* ≤ 0.001. All graphs contain n = 4-6 biological samples.

**Supplementary Figure 5:**
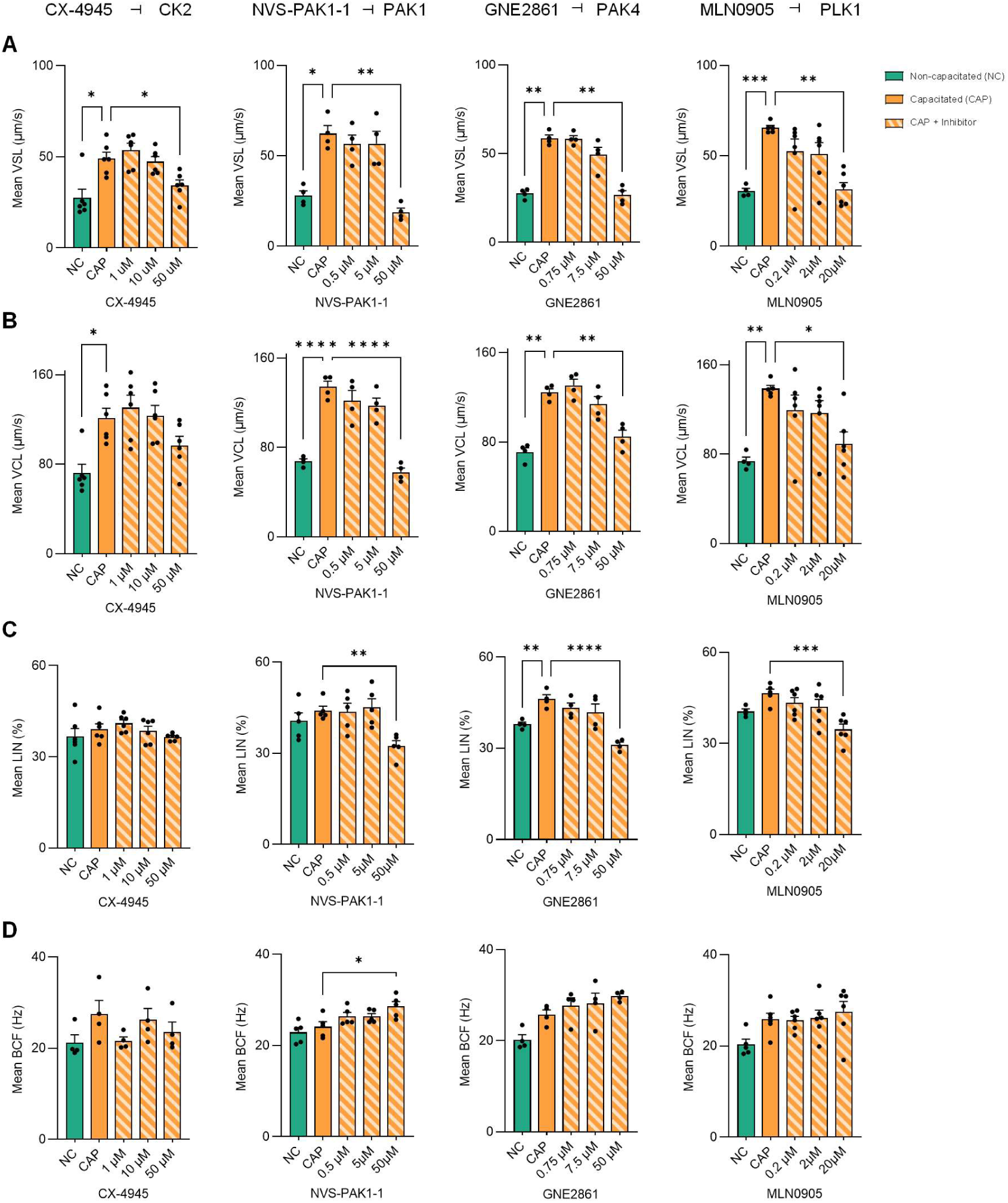
Assessment of the effect of *in vitro* kinase inhibition on sperm motility parameters. Sperm suspensions in non-capacitating BWW were “spiked” with kinase inhibitors to relevant concentrations and pre-incubated for 30 minutes before non-capacitated (NC; green) results were recorded. Samples were subsequently centrifuged and resuspended in capacitating BWW with relevant DMSO vehicle (CAP; orange) or kinase inhibitor concentrations (orange striped), incubated for 3 hours (37°C, 5% CO_2_), and subjected to motility assessment (CASA); parameters are (A) straight-line velocity (VSL), (B) curvilinear velocity (VCL), (C) linearity (LIN), and (D) beat-cross frequency (BCF). Data were subject to a non-parametric one-way ANOVA with Dunnett’s multiple comparisons test; * *p* ≤ 0.05, ** *p* ≤ 0.01, *** *p* ≤ 0.001, **** *p* ≤ 0.0001.

**Supplementary Figure 6:**
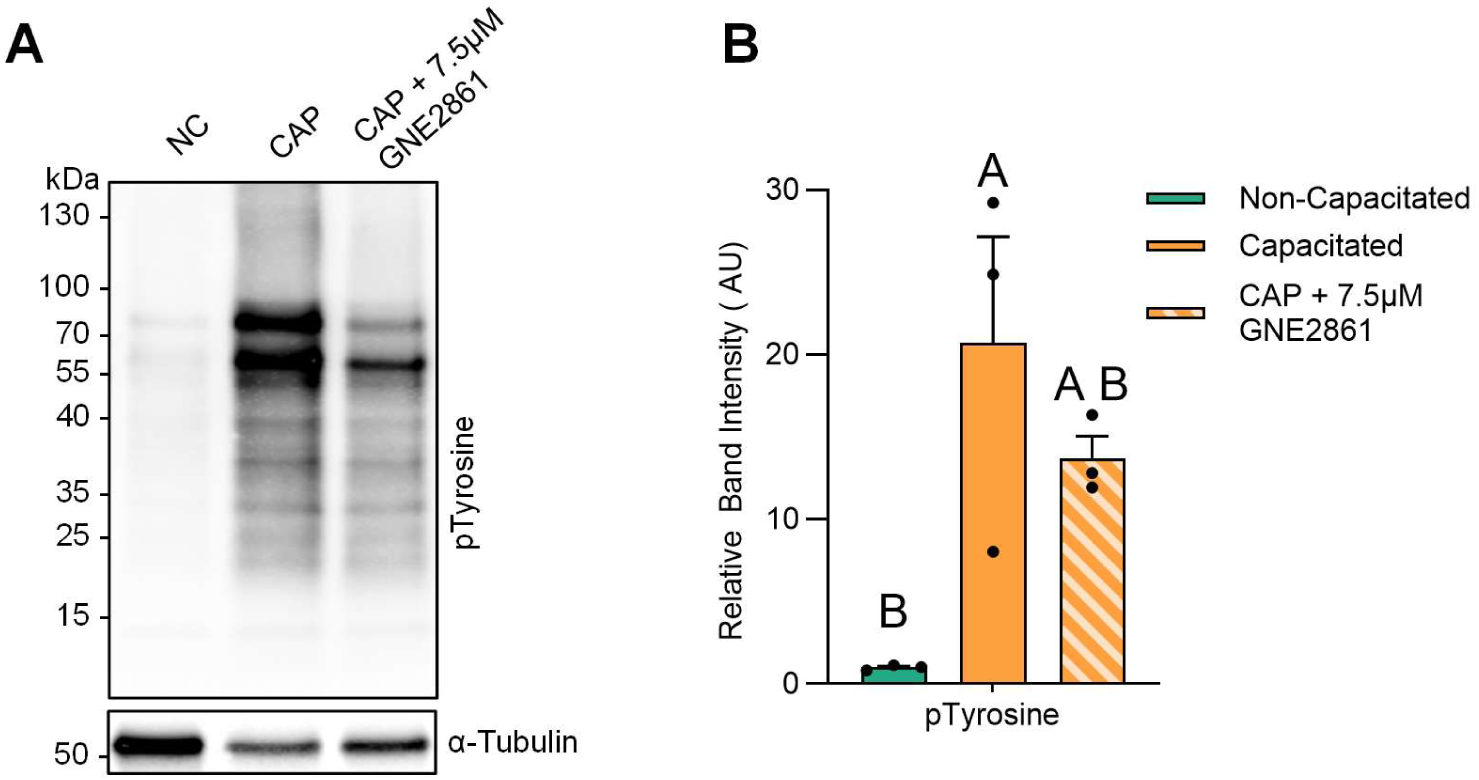
Assessment of the effect of *in vitro* kinase inhibition on sperm capacitation ability. Sperm were incubated (3 hours 37°C, 5% CO_2_) in capacitating BWW with relevant DMSO vehicle (CAP; orange) or kinase inhibitor concentrations (orange striped), and snap frozen along-side non-capacitated sperm (NC; green). To assess capacitation status samples were subjected to (A) SDS-PAGE and immunoblotting for anti-phosphotyrosine (PT66) and subsequent densitometric assessment (B). Data were subject to one-way ANOVA with Tukey’s multiple comparisons test. All graphs contain n = 3 biological samples.

**Supplementary Figure 7:**
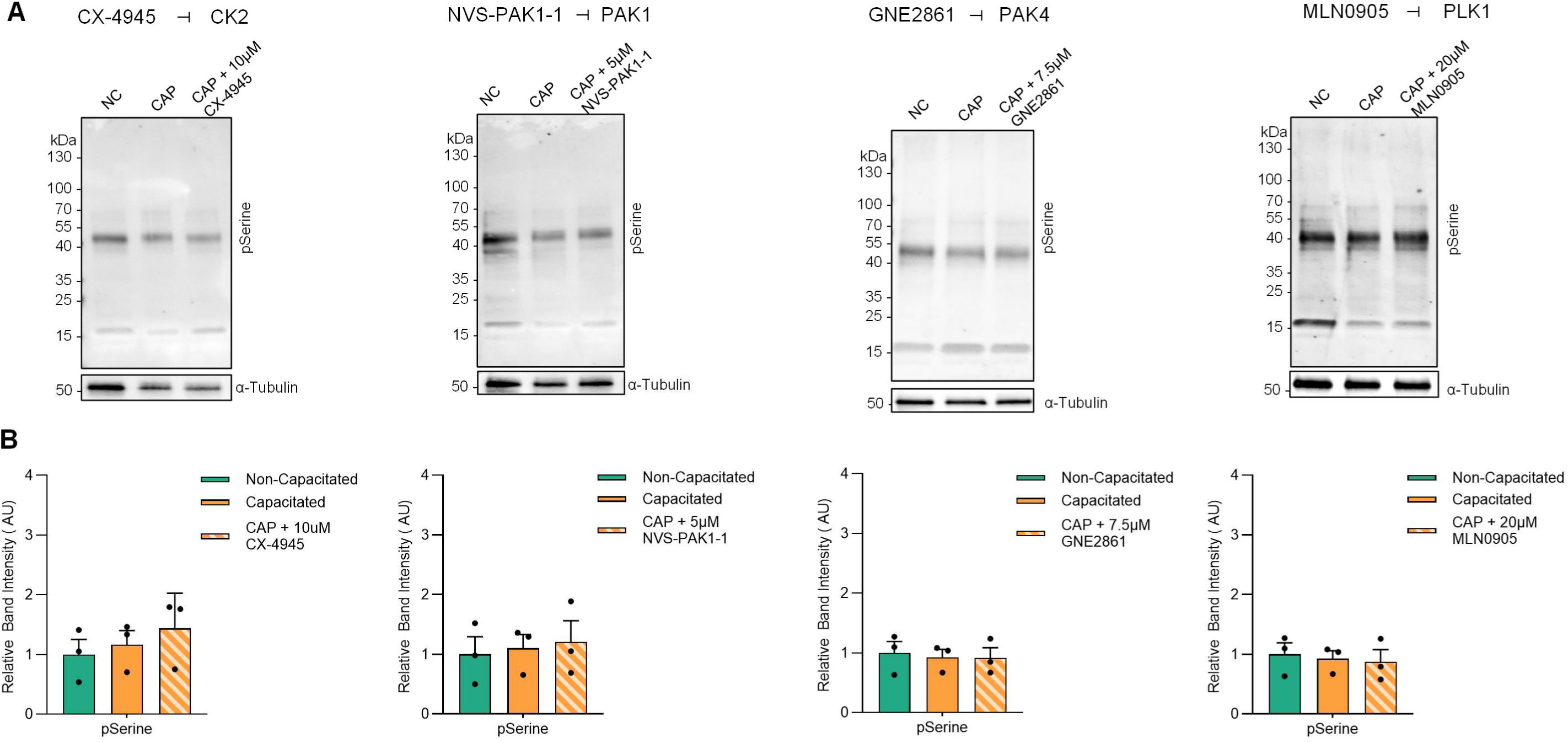
Assessment of the effect of *in vitro* kinase inhibition on sperm capacitation ability. Sperm were incubated (3 hours 37°C, 5% CO_2_) in capacitating BWW with relevant DMSO vehicle (CAP; orange) or kinase inhibitor concentrations (orange striped), and snap frozen along-side non-capacitated sperm (NC; green). Samples were subjected to SDS-PAGE and immunoblotting for (A) anti-phosphoserine (pS) and subsequent densitometric assessment (B). Data were subject to one-way ANOVA with Tukey’s multiple comparisons test; * *p* ≤ 0.05. All graphs contain n = 3 biological samples.

**Supplementary Figure 8:**
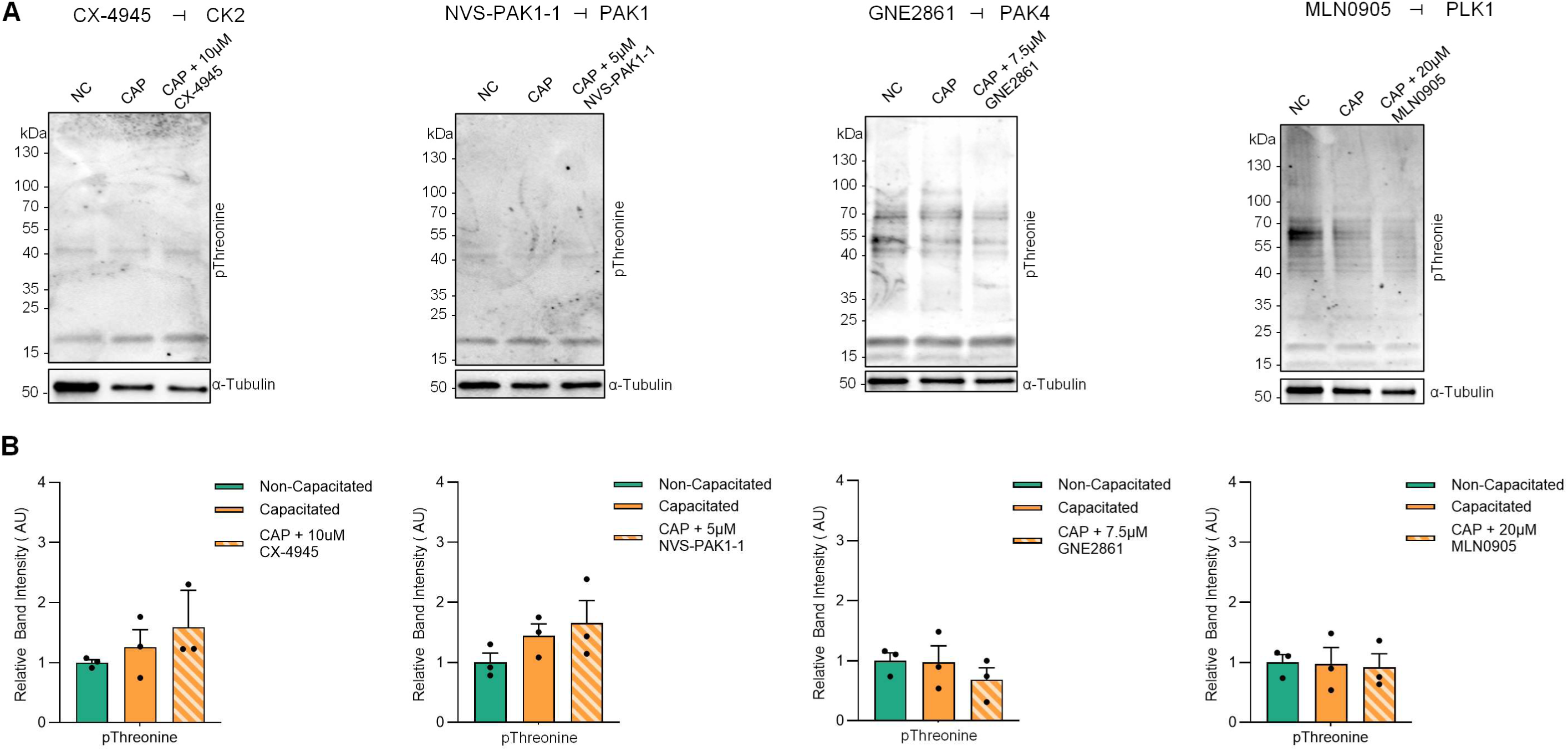
Assessment of the effect of *in vitro* kinase inhibition on sperm capacitation ability. Sperm were incubated (3 hours 37°C, 5% CO_2_) in capacitating BWW with relevant DMSO vehicle (CAP; orange) or kinase inhibitor concentrations (orange striped), and snap frozen along-side non-capacitated sperm (NC; green). Samples were subjected to SDS-PAGE and immunoblotting for (A) anti-phosphothreonine (pT) and subsequent densitometric assessment (B). Data were subject to one-way ANOVA with Tukey’s multiple comparisons test; * *p* ≤ 0.05. All graphs contain n = 3 biological samples.

**Supplementary Figure 9:**
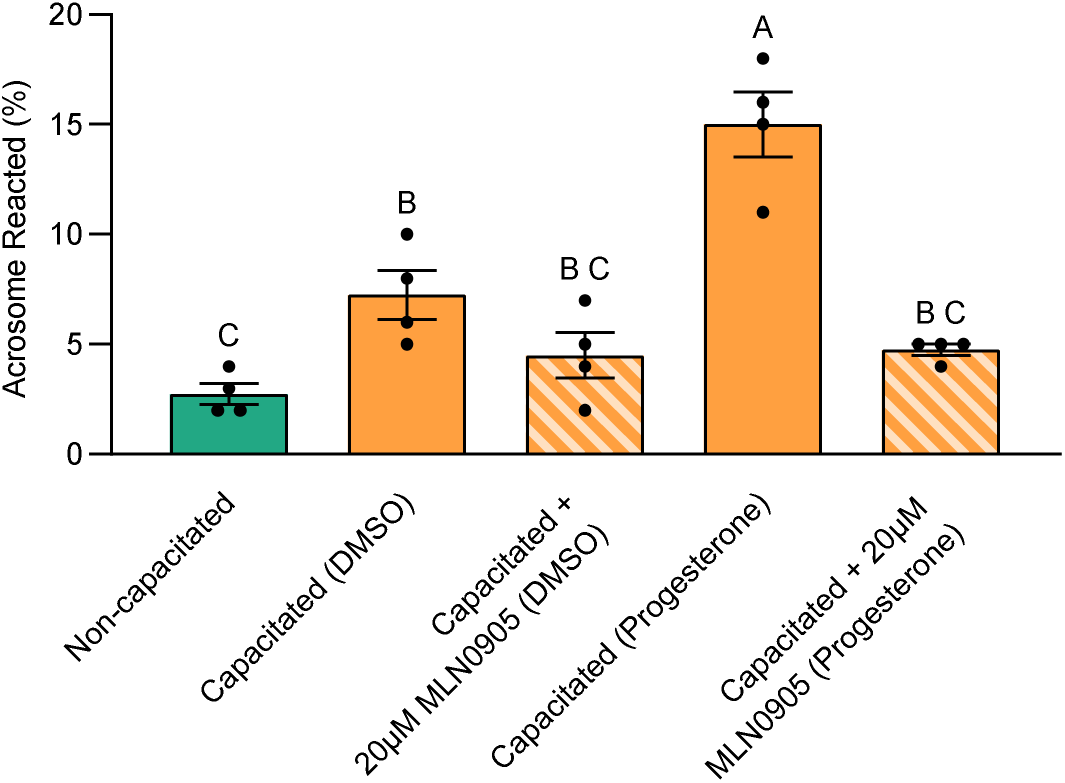
Assessment of the effect of *in vitro* kinase inhibition on sperm acrosome reaction. Sperm were incubated in capacitating BWW with a relevant a DMSO vehicle (CAP; orange) or kinase inhibitor concentrations (orange striped) for 3 hours (37°C, 5% CO_2_), and acrosome reacted along-side non-capacitated sperm. To assess acrosomal integrity cells were labelled with FITC-PSA and acrosome reacted cells counted (B). Data were subject to one-way ANOVA with Tukey’s multiple comparisons test; *p* ≤ 0.05. All graphs contain n = 4 biological samples.

**Supplementary Figure 10:**
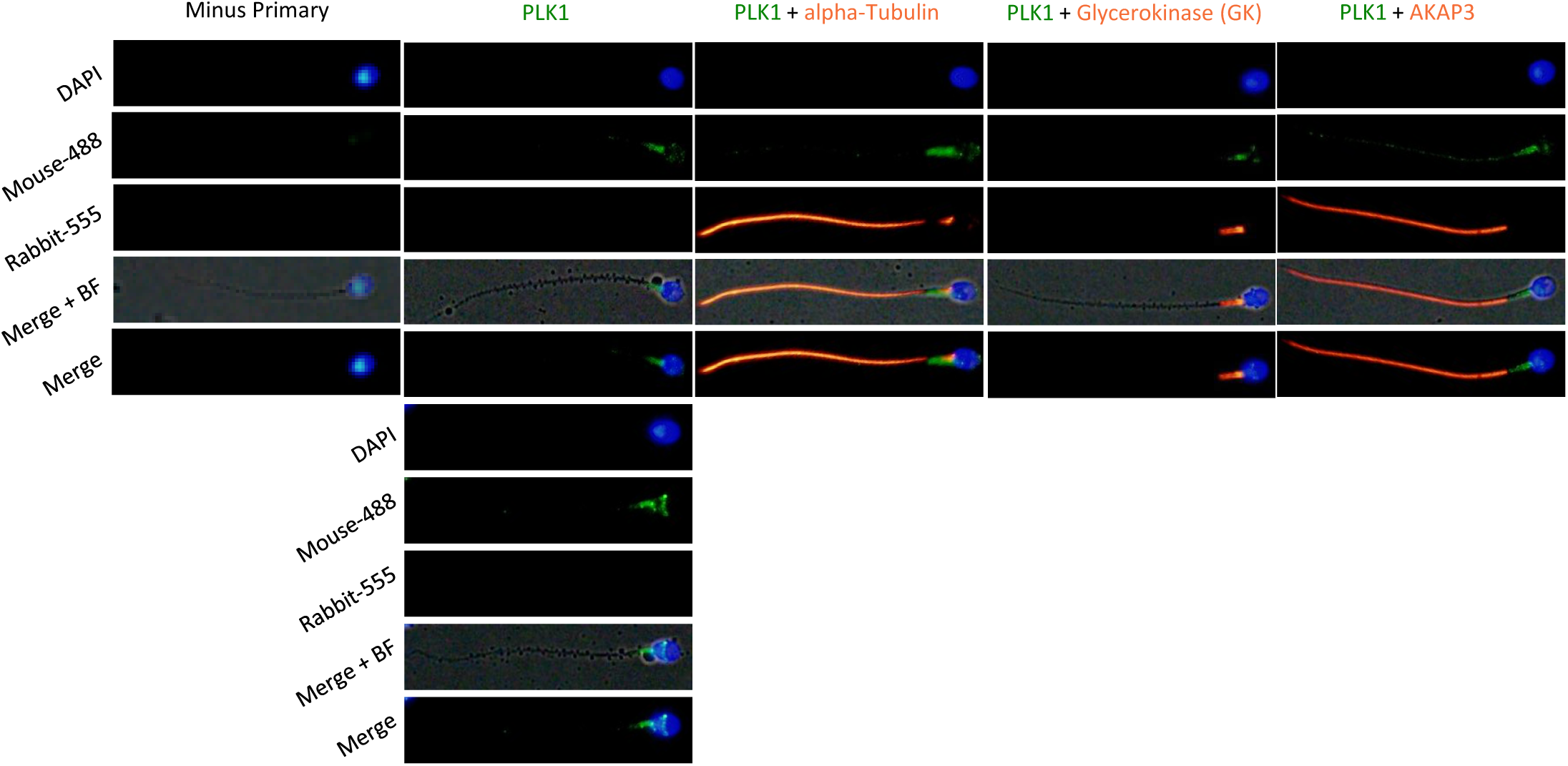
Validation of PLK1 in human sperm. Non-capacitated human sperm were immunolabelled with PLK1 (green) and co-labelled to identify localization pattern using alpha-Tubulin and AKAP3 (principle piece) and Glycerokinase (midpiece) (red). Counterstained with DAPI (Blue) and images merged with bright filed (BF).

**Supplementary Figure 11:**
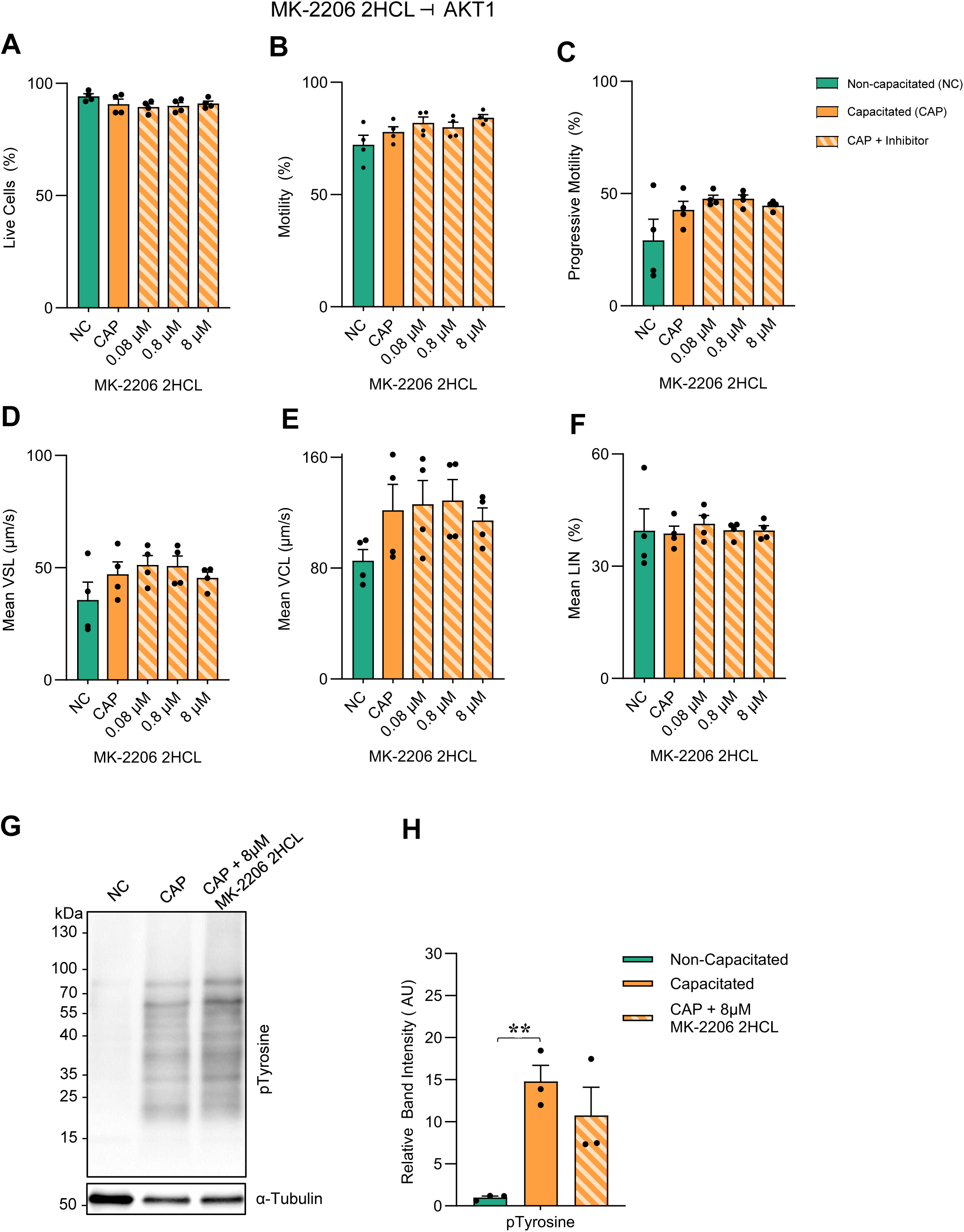
Assessment of the effect of *in vitro* AKT1 kinase inhibition using MK-2206 2HCL. Sperm suspensions in non-capacitating BWW were “spiked” with kinase inhibitors to relevant concentrations and pre-incubated for 30 minutes before non-capacitated (NC; green) results were recorded. Samples were subsequently centrifuged and resuspended in capacitating BWW with relevant DMSO vehicle (CAP; orange) or kinase inhibitor concentrations (orange striped), incubated for 3 hours (37°C, 5% CO_2_), and subjected to (A) viability (eosin) and (B-F) motility assessment (CASA). To assess capacitation status, snap frozen samples were subjected to (G) SDS-PAGE and immunoblotting for anti-phosphotyrosine (PT66) and subsequent densitometric assessment (H). Data were subject to one-way ANOVA with Tukey’s multiple comparisons test; * *p* ≤ 0.05. All graphs contain n = 3-4 biological samples.

**Table S1:** Human sperm phosphoproteome.

**Table S2:** Ingenuity Pathway Analysis Biological Processes.

**Table S3:** Putative kinases predicted by Phosphomatics *in silico* analysis Table S4: Semen analysis of sperm donors

**Table S5:** Antibody table.

